# Changes in Walking Function and Neural Control following Pelvic Cancer Surgery with Reconstruction

**DOI:** 10.1101/2024.02.27.582423

**Authors:** Geng Li, Di Ao, Marleny M. Vega, Payam Zandiyeh, Shuo-Hsiu Chang, A.N. Penny, Valerae O. Lewis, Benjamin J. Fregly

**Affiliations:** Rice Computational Neuromechanics Laboratory, Department of Mechanical Engineering, Rice University, Houston, TX, USA; Biomotion Laboratory, Department of Orthopedic Surgery, McGovern Medical School at the University of Texas Health Science Center at Houston, Houston, TX, USA; Department of Physical Medicine and Rehabilitation, McGovern Medical School at the University of Texas Health Science Center at Houston, Houston, TX, USA; Department of Orthopedic Oncology, University of Texas MD Anderson Cancer Center, Houston, TX, USA

**Keywords:** pelvic sarcoma, custom implant, orthopedic oncology, instrumented gait analysis, walking biomechanics, muscle synergies, neuromusculoskeletal modeling

## Abstract

Surgical planning and custom prosthesis design for pelvic cancer patients is challenging due to the unique clinical characteristics of each patient and the significant amount of pelvic bone and hip musculature often removed. Limb-sparing internal hemipelvectomy surgery with custom prosthesis reconstruction has become a viable option for this patient population. However, little is known about how post-surgery walking function and neural control change from pre-surgery conditions. This case study combined comprehensive human movement data collection with personalized neuromusculoskeletal computer modeling to provide a thorough assessment of pre-to post-surgery changes in walking function and neural control for a single pelvic sarcoma patient who received internal hemipelvectomy surgery with custom prosthesis reconstruction. Extensive walking data (video motion capture, ground reaction, and EMG) were collected from the patient before surgery and after plateau in recovery after surgery. Pre- and post-surgery personalized neuromusculoskeletal computer models of the patient were then constructed using the patient’s pre- and post-surgery walking data. These models were used to calculate the patient’s pre- and post-surgery joint motions, joint moments, and muscle synergies. The calculated muscle synergies were described by time-invariant synergy vectors and time-varying synergy activations, were consistent with the patient’s experimental EMG data, and produced the patient’s experimental joint moments found via inverse dynamics. The patient’s post-surgery walking function was marked by a slower self-selected walking speed coupled with several compensatory mechanisms necessitated by lost or impaired hip muscle function, while the patient’s post-surgery neural control demonstrated a dramatic change in coordination strategy (as evidenced by modified synergy vectors) with little change in recruitment timing (as evidenced by conserved synergy activations). Furthermore, the patient’s post-surgery muscle activations were fitted accurately using the patient’s pre-surgery synergy activations but poorly using the patient’s pre-surgery synergy vectors. These results provide valuable information about which aspects of post-surgery walking function could potentially be improved through modifications to surgical decisions, custom prosthesis design, or rehabilitation protocol, as well as how computational simulations could be formulated to predict post-surgery walking function reliably given a patient’s pre-surgery walking data and the planned surgical decisions and custom prosthesis design.

## 1 Introduction

Pelvic sarcomas account for up to 20% of the approximately 2,500 primary bone tumor cases reported in the United States each year, where most patients are 25 years of age or younger (Mayerson et al., 2014, Morris, 2010). The complex anatomy of the pelvic region and the heterogeneity across patients in how tumors infiltrate the pelvis make surgical treatment for pelvic sarcomas challenging. Thanks to recent advances in medical imaging and multimodal oncological treatments (Puchner et al., 2017), internal hemipelvectomy surgery (Bickels and Malawer, 2001) has gained wider use for removing bone and soft tissues infiltrated by the tumor while sparing the limb. In pelvic sarcoma cases where the hip joint is also infiltrated, the acetabulum and femoral head must also be removed. Initially, no reconstruction of the hip joint was the primary surgical option, which produces functional walking ability but with an abnormal gait pattern, some functional limitations, and a long recovery time, along with a risk of developing low back pain and scoliosis due to a significant limb-length discrepancy (Lackman et al., 2009; Wedemeyer and Kauther, 2011). More recently, custom prosthesis reconstruction with a total hip replacement has become a viable surgical option, reducing functional limitations and recovery time while eliminating the limb-length discrepancy, thereby also reducing the risk of developing low back pain and scoliosis (Lewis, 2014; Chao et al., 2015).

Unfortunately, few experimental studies have assessed how well custom prosthesis reconstruction with a total hip replacement is able to restore pre-surgery walking function and neural control. One study reported peak vertical ground reaction forces post-surgery to quantify the effects of an external hip stabilizing device on gait symmetry (Akiyama et al., 2016). A second study measured joint motion and metabolic energy expenditure to assess walking function following a two-year rehabilitation program (Wingrave and Jarvis, 2019). A third study collected comprehensive post-surgery gait data including vertical ground reaction forces, joint motions, joint moments, and electromyography (EMG) data to assess motor performance and walking (Valente et al., 2022). In all three of these studies, operated side biomechanical quantities were compared to non-operated side quantities following surgery, since no pre-surgery data were available from the same patients for comparison. Furthermore, while two studies reported extensive side-to-side comparisons of joint motion data, no study has presented ground reaction data for all three components of ground reaction force or joint moment data for all lower body joints. In addition, no study to date has quantified how a patient’s neural control strategy changes in response to the surgery. Since the surgery involves removal of multiple hip muscles, one might expect that a significant change in neural control would be required to allow a patient to walk following such extensive surgery. Even if comprehensive experimental walking data were available before and after surgery from the same patient to quantify changes in walking function, such data alone would not provide an objective means for improving surgical planning or custom pelvic prosthesis design.

Coupling comprehensive human movement data collection with personalized neuromusculo-skeletal computer modeling provides a novel approach for assessing changes in not only biomechanical quantities but also neural control while also providing an objective means for predicting how planned surgical decisions will affect post-surgery walking function. While a comprehensive data set is needed to personalize a neuromusculoskeletal model (Meyer et al., 2016; Arones et al., 2020), the personalized model can then provide novel insights that would not be available from the data alone. For example, a personalized neuromusculoskeletal model developed from a patient’s movement data can be used to analyze joint contact loads in knee osteoarthritis, muscle force generation in cerebral palsy, and neural control capabilities in stroke (Fregly et al., 2012; Meyer et al., 2016; Pitto et al., 2019). In particular, EMG data can become functional (i.e., it generates the correct joint moments) and not just descriptive through the use of a personalized neuromusculoskeletal model. Several recent studies have collected comprehensive human movement data sets from individuals who were implanted with instrumented total knee replacements (Taylor et al., 2004; Fregly et al., 2012) or who suffered a stroke (Meyer et al., 2016, 2017; Sauder et al., 2019). These data sets include joint motion, ground reaction, and EMG data collected for walking and other functional tasks, and researchers have published associated personalized neuromusculoskeletal models that have been used to predict patient walking function under new conditions (Meyer et al., 2016), while receiving functional electrical stimulation (Sauder et al., 2019), while using crutches (Febrer-Nafría et al., 2021), or following pelvic cancer surgery (Vega et al., 2022).

This case study presents the most comprehensive quantitative assessment to date of walking function (joint motions, joint moments, ground reactions) and neural control (muscle synergies) changes produced by internal hemipelvectomy surgery with custom prosthesis reconstruction. The study combines collection of comprehensive pre- and post-surgery gait data (video motion capture, ground reaction, EMG) from a single patient with development of pre- and post-surgery personalized neuromusculoskeletal computer models of the same patient. The personalized models make it possible to analyze not only biomechanical changes, which provide valuable information about which aspects of post-surgery walking function remain abnormal and thus should be the targets for physical therapy, but also neural control changes, which provide valuable information on how neural control should be modeled to predict post-surgery walking function given pre-surgery walking data and the planned surgical decisions. Neural control changes are quantified using muscle synergies, which provide a low-dimensional description of how the central nervous system controls a large set of individual muscles (D’Avella et al., 2003; Tresch et al., 2006; Chvatal and Ting, 2013).

## 2 Methods

### 2.1 Experimental Data Collection

Experimental walking and CT scan data were collected pre- and post-surgery from a single patient with a pelvic sarcoma who received internal hemipelvectomy surgery with custom prosthesis reconstruction. The patient (sex: male, age: 46 years, height: 1.73 m, mass: 82.5 kg) gave written informed consent, and all data collection and subsequent computational analyses were approved by the institutional review boards of MD Anderson Cancer Center, the University of Texas Health Science Center Houston, and Rice University. The surgery involved resection of the tumor, hip joint, and surrounding muscles in the right hemipelvis in the pubic and acetabular regions, and the custom prosthesis included a total hip replacement.

Pre- and post-surgery walking and CT scan data were collected using identical protocols, where the subject walked at his self-selected speed (1.0 m/s pre-surgery and 0.5 m/s post-surgery). Experimental walking data included ground reaction, video motion capture, and EMG data. Ground reaction data were collected from a split-belt instrumented treadmill (Bertec Corp., Columbus, OH, USA) with belts tied. Video motion capture data were collected using an optical motion capture system (Qualisys AB, Gothenburg, Sweden). A total of 36 retroreflective markers were placed on the feet, legs, pelvis, torso, and arms consistent with a previous study (Meyer 2016). A static standing trial was performed to facilitate a subsequent musculoskeletal model scaling operation (see below). Walking trials at a single speed were performed to measure the subject’s gait kinematics, kinetics, and muscle activity. EMG data were collected from 15 lower extremity muscles per leg (Table 1) using both surface and fine wire electrodes (Cometa, Bareggio, Italy). EMG data processing was consistent with a previous study (Meyer et al., 2017), including high-pass filtering at 40 Hz, demeaning, rectifying, and low-pass filtering using a variable cut-off frequency dependent on the gait period (Hug, 2011). CT scan data of the subject’s entire pelvis and both proximal femurs were collected to facilitate the creation of pre- and post-surgery musculoskeletal models with personalized geometry. In addition, a geometric model of the subject’s custom pelvic prosthesis was made available by the orthopedic implant manufacturer (Onkos Surgical, Parsippany, NJ, USA).

**Table 1:** List of lower extremity muscles in the musculoskeletal model, the source from which the muscle excitations were acquired and the acquisition method. The shaded muscles were surgically removed from the operated (right) leg during surgery.

| Muscle Names (Abbreviation) | Muscle Excitation Source | Muscle Excitation Acquisition Method |
| --- | --- | --- |
| adductor brevis (ADB)<br>adductor longus (ADL)<br>adductor magnus distal (ADM1)<br>adductor magnus ischial (ADM2)<br>adductor magnus middle (ADM3)<br>adductor magnus proximal (ADM4)<br>gracilis (GRAC) | adductor longus | Measured |
| biceps femoris long head (BFLH)<br>biceps femoris short head (BFSH) | biceps femoris long head |  |
| gastrocnemius lateral (GL) | gastrocnemius lateral |  |
| gastrocnemius medial (GM) | gastrocnemius medial |  |
| gluteus maximus superior (GMA1)<br>gluteus maximus middle (GMA2)<br>gluteus maximus inferior (GMA3) | gluteus maximus |  |
| gluteus medius anterior (GME1)<br>gluteus medius middle (GME2)<br>gluteus medius posterior (GME3)<br>gluteus minimus anterior (GMI1)<br>gluteus minimus middle (GMI2)<br>gluteus minimus posterior (GMI3) | gluteus medius |  |
| iliacus (IL)<br>psoas major superior (PS1)<br>psoas major middle (PS2)<br>psoas major inferior (PS3) | iliopsoas † |  |
| peroneus brevis (PB)<br>peroneus longus (PL) | peroneus |  |
| rectus femoris (RF)<br>pectineus (PECT) | rectus femoris |  |
| semimembranosus (SM)<br>semitendinosus (ST) | semimembranosus |  |
| vastus medialis (VM)<br>vastus intermedius (VI) | vastus medius |  |
| vastus lateralis (VL) | vastus lateralis |  |
| soleus (SOL) | soleus |  |
| tibialis anterior (TA) | tibialis anterior |  |
| tibialis posterior (TP) | tibialis posterior |  |
| extensor digitorum longus (EDL)<br>extensor hallucis longus (EHL) | extensor digitorum longus | Synergy Extrapolation |
| flexor digitorum longus (FDL)<br>flexor hallucis longus (FHL) | flexor digitorum longus |  |
| gemellus (GEM)<br>piriformis (PIRI)<br>quadratus femoris (QF) | piriformis |  |
| sartorius (SART) | sartorius |  |
| tensor fasciae latae (TFL) | tensor fasciae latae |  |

739 † EMG measurement of the right iliopsoas was unavailable pre- and post-surgery.

### 2.2 Neuromusculoskeletal Model Analyses

Personalized neuromusculoskeletal computer models of the subject were constructed to represent pre- and post-surgery conditions, where all geometric musculoskeletal modeling was performed in OpenSim (Delp et al., 2007; Seth et al., 2018). The model personalization process started with a generic OpenSim model constructed by combining previously published lower extremity models (Arnold et al., 2010; Rajagopal et al., 2016; Lai et al., 2017) and lumbar-spine models (Christophy et al., 2012; Bruno et al., 2015). This process was performed to produce a new generic model that possessed all of the muscles required for the present study. The resulting generic model possessed 58 trunk muscles and 45 muscles in each leg to actuate the following rotational joints with their associated number of degrees of freedom: hip (3), knee (1), ankle (1), subtalar (1), and toe (1) on each leg along with lumbosacral (3).

Starting from this generic model, we constructed a pre-surgery model of the patient using his pre-surgery walking and CT scan data as described in a previous study (Li et al., 2022). In brief, the body segments in the generic model were scaled to match the subject using static trial motion capture data and the OpenSim Model Scaling Tool. The one exception was the pelvis, whose dimensions were scaled separately in all three directions to match the subject’s pelvis geometry as determined from pre-surgery CT scan data. The scaled generic pelvis geometry was then replaced with personalized pelvis geometry constructed from pre-surgery CT scan data. Pelvis muscle attachment locations were adjusted to be on bony anatomy by following a codified workflow (Modenese et al., 2018) using nmsBuilder (Valente et al., 2017).

Starting from the pre-surgery model, we constructed a post-surgery model of the patient by modifying his pre-surgery model to account for implantation of a custom pelvic prosthesis and the surgical decisions made by the orthopedic oncologist. To account for implantation of a custom pelvic prosthesis, we developed a bone-prosthesis geometric model of the patient’s pelvis and operated femur. For the pelvis with custom prosthesis, we first used a two-step process of global registration and fine alignment in Geomagic Wrap (3D Systems, Morrisville, NC, USA) to align a post-surgery geometric model of the pelvis plus custom prosthesis with the pre-surgery geometric model of the pelvis. The pre-surgery pelvis geometric model was subsequently replaced by the post-surgery pelvis-prosthesis geometric model (Fig. 1A). The hip joint center on the operated side was updated with the center of a sphere used to fit the inner surface of the acetabular component (Fig. 1B). For the post-surgery femur with femoral component, we obtained the bone and implant geometry from post-surgery CT scan data. The femoral geometry in the pre-surgery model on the operated side was then re-scaled only in the superior-inferior direction to account for the slight change in femur length due to implantation of the femoral component. After the post-surgery femur with femoral component geometry was aligned to the re-scaled pre-surgery femoral geometry using Geomagic Wrap, the re-scaled pre-surgery femur geometry was replaced with the post-surgery femur with femoral component geometry. To account for the surgical decisions made by the orthopedic oncologist, we set to zero the peak isometric strength of 15 muscle heads surgically removed from the operated leg (Supplementary Table 1). For muscles that were surgically detached and reattached, the pre-surgery peak isometric strength and geometry were retained in the post-surgery model.

**Figure 1.**
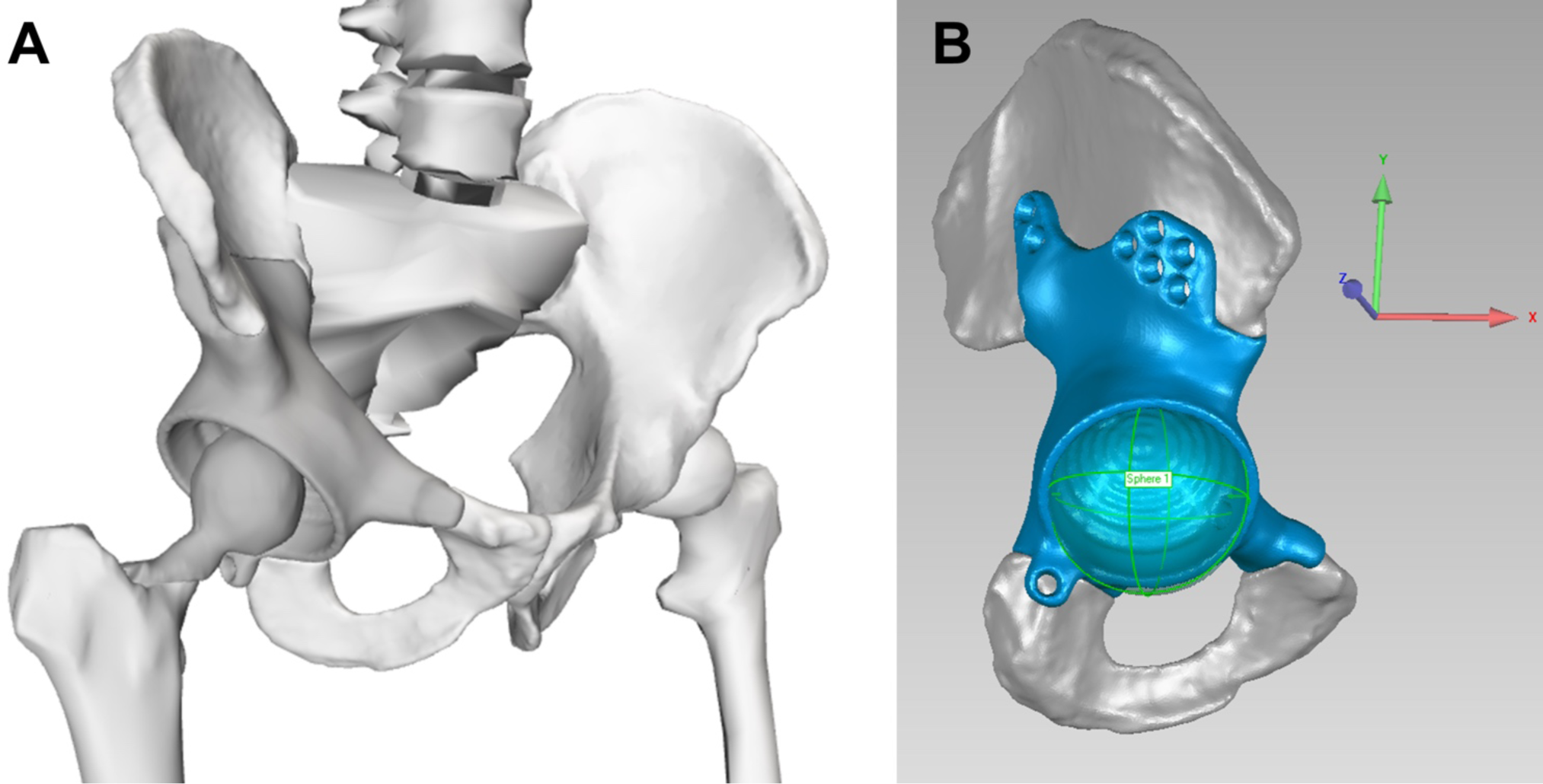
A. Geometric model of the remaining pelvic bone, custom prosthesis, and total hip replacement. B. Updated hip joint center on the operated side, as determined from the center of the sphere used to fit the inner surface of the acetabular component.

A sequence of three standard OpenSim operations were performed using the patient’s pre- and post-surgery musculoskeletal models to quantify changes in gait biomechanics and to generate input data needed for EMG-driven model calibration (Lloyd and Besier, 2003; Buchanan et al., 2004; Sartori et al., 2014). The OpenSim Inverse Kinematics Tool was used to compute joint kinematics using the patient’s motion capture marker data. The OpenSim Inverse Dynamics Tool was used to compute joint moments using the patient’s calculated joint kinematics and experimental ground reaction data. The OpenSim Muscle Analysis Tool was used to compute muscle-tendon lengths and muscle moment arms using the patient’s joint kinematics. Ten pre- and post-surgery gait cycles were identified that possessed the smallest root-mean-square error (RMSE) values for joint angle and joint moment curves with respect to corresponding mean curves. Data from these gait cycles were used for all subsequent analyses.

An EMG-driven neuromusculoskeletal model was calibrated for the hip, knee, ankle, and subtalar joints of each leg to estimate the patient’s lower extremity muscle-tendon model parameter values, muscle excitations, and muscle activations before and after surgery (Meyer et al., 2017; Ao et al., 2020, 2022, 2023). Muscle excitations were derived from processed EMG data while muscle activations were produced by applying electromechanical delays and activation dynamics to the muscle excitations. Muscle excitations included experimental and residual excitations for muscles with associated EMG data and predicted excitations for muscles without associated EMG data (Table 1). Experimental excitations were normalized using EMG scale factor design variables found during the EMG-driven model calibration process. Residual excitations were small changes applied to experimental excitations that enabled more accurate estimation of predicted excitations (Ao et al., 2022). Residual and predicted muscle excitations were estimated using synergy extrapolation (Bianco et al., 2018; Ao et al., 2020, 2022), which applied principal component analysis to the patient’s experimental excitations to find experimental synergy excitations. These synergy excitations were multiplied by synergy vector weight design variables found during the EMG-driven model calibration process to produce the estimated residual and predicted excitations.

A multi-objective optimization problem was solved to perform the EMG-driven model calibration process with synergy extrapolation for each leg before and after surgery (Bianco et al., 2018; Ao et al., 2020, 2022). The design variables adjusted by the optimization were the following: 1. Activation dynamics model parameters: EMG scale factors, electromechanical delays, activation time constants, and activation non-linear shape factors (He et al., 1991; Lloyd and Besier, 2003), 2. Hill-type muscle-tendon model parameters: optimal fiber length (*l*_*Mo*_) and tendon slack length (*l*_*Ts*_) (Zajac, 1989), and 3. Synergy vector weights for constructing residual and predicted muscle excitations. Peak isometric strength parameter values for all lower extremity muscle-tendon models were calculated using published regression relationships derived from MRI data (Handsfield et al., 2014). Muscle-tendon model parameter values for muscles spanning the lumbosacral joint were taken from a previously published EMG-driven modeling study that used the same pre-surgery walking data collected from the same patient (Li et al., 2022).The primary cost function term minimized the sum of squares of differences between experimental joint moments calculated via inverse dynamics and model joint moments calculated using the patient’s personalized neuromusculoskeletal model. Secondary cost function terms minimized the sum of squares of residual and predicted muscle excitations. The EMG-driven model calibration process was first performed for each leg using pre-surgery walking data, and the resulting parameter values were used as the initial guess when the calibration process was repeated using post-surgery walking data. Each calibrated EMG-driven model yielded activations dynamics and muscle-tendon model parameter values along with estimates of muscle excitations and activations, henceforth referred to collectively as “muscle controls.”

### 2.3 Lower Extremity Muscle Synergy Analyses

The pre- and post-surgery muscle controls for each leg were decomposed into a lower dimensional space using muscle synergy analysis (MSA). MSA was performed via non-negative matrix factorization (NMF) (Tresch et al., 2006; Rabbi et al., 2020) using the MATLAB ‘nnmf’ function (MathWorks, Natick, MA, USA) on both muscle excitations and muscle activations for each pre- and post-surgery gait cycle using only muscles with experimental EMG data (Table 1). Each muscle synergy consisted of a time-varying synergy control along with a corresponding time-invariant synergy vector containing muscle-specific weights. Synergy controls provided information about recruitment timing (i.e., when groups of muscles were co-activated) while synergy vectors provided information about coordination strategy (i.e., how groups muscles were co-activated). Synergy controls extracted from muscle excitations were referred to as synergy excitations, while those extracted from muscle activations were referred as synergy activations. MSA was performed assuming 4, 5, 6 and 7 synergies for each leg as this range covers the typical number of synergies required to adequately represent muscle activity during gait (Ivanenko et al., 2004; Meyer et al., 2016). The variability accounted for (VAF) metric was used to quantify how well the calculated muscle synergies could represent the original muscle controls. Since muscle synergies were calculated for each gait cycle separately and the NMF algorithm does not output muscle synergies in any particular order, muscle synergies were sorted based on cosine similarity between synergy vectors across gait cycles (Banks et al., 2017).

To account for how a difference between pre- and post-surgery walking speed likely affected the magnitude of the synergy controls (Clark et al., 2010), we introduced a new magnitude quantity into the muscle synergy decomposition equation (Eq. 1).

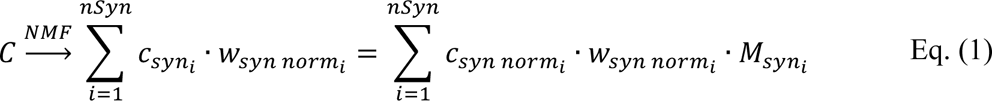

where *C* are muscle controls while *c*_*syn*_ and *w*_*syn norm*_ are synergy controls and normalized synergy vectors, respectively. Synergy vectors are normalized automatically to a magnitude of one by the Matlab ‘nnmf’ algorithm, but synergy controls are not normalized. Consequently, we normalized each synergy control to a maximum value of one and added a new parameter *M*_*syn*_to represent the magnitude of the synergy. With this modification, synergy controls provide information about only recruitment timing, synergy vectors information about only coordination strategy, and synergy magnitudes information about only synergy magnitude, making it easier to quantify synergy changes produced by changes in walking speed.

Since all muscle synergies were calculated using muscle excitations and activations produced by a calibrated EMG-driven lower extremity model, the resulting muscle synergies where functional rather than merely descriptive. That is, when calculated muscle synergies were input into the appropriate EMG-driven leg model, they produced hip, knee, ankle, and subtalar joint moments that closely matched the patient’s experimental joint moments calculated via inverse dynamics, with the closeness of the match determined by the VAF of the calculated synergies. Standard MSA is applied to EMG data alone and will not produce the correct lower extremity joint moments when the calculated synergies are input into a lower extremity neuromusculoskeletal model.

### 2.4 Post-surgery Muscle Control Reconstruction

We explored three options for how post-surgery muscle excitations and activations could be reconstructed using corresponding pre-surgery muscle synergy information. For the first option, termed the Fixed Synergy Vector method, the post-surgery synergy vectors were assumed to be identical to the pre-surgery synergy vectors, implying the coordination strategy was conserved, and the corresponding post-surgery synergy controls needed to reconstruct the post-surgery muscle controls were calculated. For the second option, termed the Fixed Synergy Control method, the post-surgery synergy controls were assumed to be identical to the pre-surgery synergy controls, implying the recruitment timing was conserved, and the corresponding post-surgery synergy vectors required to reconstruct the post-surgery muscle controls were calculated. For the third option, termed the Shifted Synergy Control method, the post-surgery synergy controls were assumed to be identical to the pre-surgery synergy controls except with small time shifts, implying the recruitment timing was conserved apart from small time shifts, and the corresponding post-surgery synergy vectors required to reconstruct the post-surgery muscle controls were calculated.

We implemented all three options for reconstructing post-surgery muscle controls using the MSA results obtained for the pre-surgery muscle controls. For the Fixed Synergy Vector method, the mean synergy vectors across all pre-surgery gait cycles were defined as the fixed synergy vectors, and the synergy controls required to reconstruct the post-surgery muscle controls for each gait cycle were found (Eq. (2)). For the Fixed Synergy Control method, the mean synergy controls across all pre-surgery gait cycles were defined as the fixed synergy controls, and the synergy vectors required to reconstruct the post-surgery muscle controls for each gait cycle were found (Eq. (3)). For the Shifted Synergy Control method, each pre-surgery synergy control was shifted in time to maximize its cosine similarity with the corresponding post-surgery synergy control (Supplementary Fig. 6). The shifted pre-surgery synergy controls were then used as the fixed synergy controls in Eq. (3), and the synergy vectors required to reconstruct the post-surgery muscle controls for each gait cycle were found. Synergy controls and vectors were considered to be non-negative and therefore nonnegative linear least-squares optimization problems were solved using the MATLAB ‘lsqnonneg’ function. To calculate synergy controls for the Fixed Synergy Vector method, we used the following problem formulation:

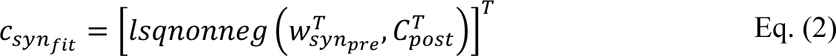

where *c*_*synfit*_ are synergy controls for each gait cycle that produce the best fit of *C*_*post*_, the post-surgery muscle controls for each gait cycle, using *w*_*synpre*_, the mean pre-surgery synergy vectors. Similarly, to calculate synergy vectors for the Fixed or Shifted Synergy Control method, we solved:

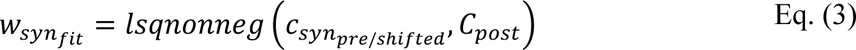

where *w*_*synfit*_ are the synergy vectors for all gait cycles that produce the best fit of *C*_*post*_, the post-surgery muscle controls for each gait cycle, using *c*_*synshifted*_, the mean pre-surgery synergy controls without and with time shifting.

To evaluate how well the post-surgery muscle controls were reconstructed, we calculated the variability accounted for (VAF) between reconstructed and experimental muscle controls for each post-surgery gait cycle. The evaluation was performed for each combination of the following three methodological choices: 1) selected reconstruction option (Fixed Synergy Vector, Fixed Synergy Control, or Shifted Synergy Control method), 2) type of muscle controls to be reconstructed (excitations or activations), and 3) number of synergies used per leg (4, 5, 6, or 7).

### 2.5 Statistical Analyses

Statistical analyses were used to compare time-varying pre- and post-surgery biomechanical and neural control quantities and time-invariant VAF values for reconstructing post-surgery muscle controls from pre-surgery muscle synergies produced by different combinations of methodological choices. To compare time-varying quantities (ground reaction forces, joint angles, joint moments, and synergy controls) before and after surgery, we used statistical parametric mapping (SPM) (Penny et al., 2011), which presents statistical results in the same temporal space as the original data, making interpretation of results straightforward (Pataky, 2012). SPM used two-tailed two sample t-tests (*p* < 0.05) to compare pre- and post-surgery time-varying quantities across the gait cycle. SPM analyses were implemented using the ‘ttest2’ function from the opensource SPM code ‘spm1d’ in MATLAB (Pataky, 2012). To compare time-invariant VAF values produced by different methodological choices for reconstructing post-surgery muscle controls from pre-surgery muscle synergies, we used paired sample t-tests (*p* < 0.05) implemented using the MATLAB ‘ttest’ function. Each sample consisted of ten VAF values, one for each post-surgery gait cycle, calculated using a specified methodological choice (e.g., Fixed Synergy Vector method applied to muscle excitations using 4 synergies).

## 3 Results

### 3.1 Walking Function Changes

Significant differences were observed between the pre- and post-surgery ground reaction force (GRF), joint motion, and joint moment data obtained from the patient’s experimental walking data and personalized neuromusculoskeletal models. For ground reaction data (Fig. 2), the vertical GRF during the loading phase and mid-stance were different for the operated and non-operated legs between post and pre-surgery conditions, as was the vertical GRF during the unloading phase for the operated leg (*p* < 0.001). The decrease in vertical GRF during operated leg unloading and the increase in vertical GRF during non-operated leg loading each started earlier in the gait cycle during post-surgery walking, suggesting that the non-operated leg was compensating for the operated leg. Propulsive force shortly before toe-off and braking force following heel strike were also significantly lower post-surgery than pre-surgery for both legs (*p* < 0.001), possibly related to the decrease in self-selected walking speed post-surgery.

**Figure 2.**
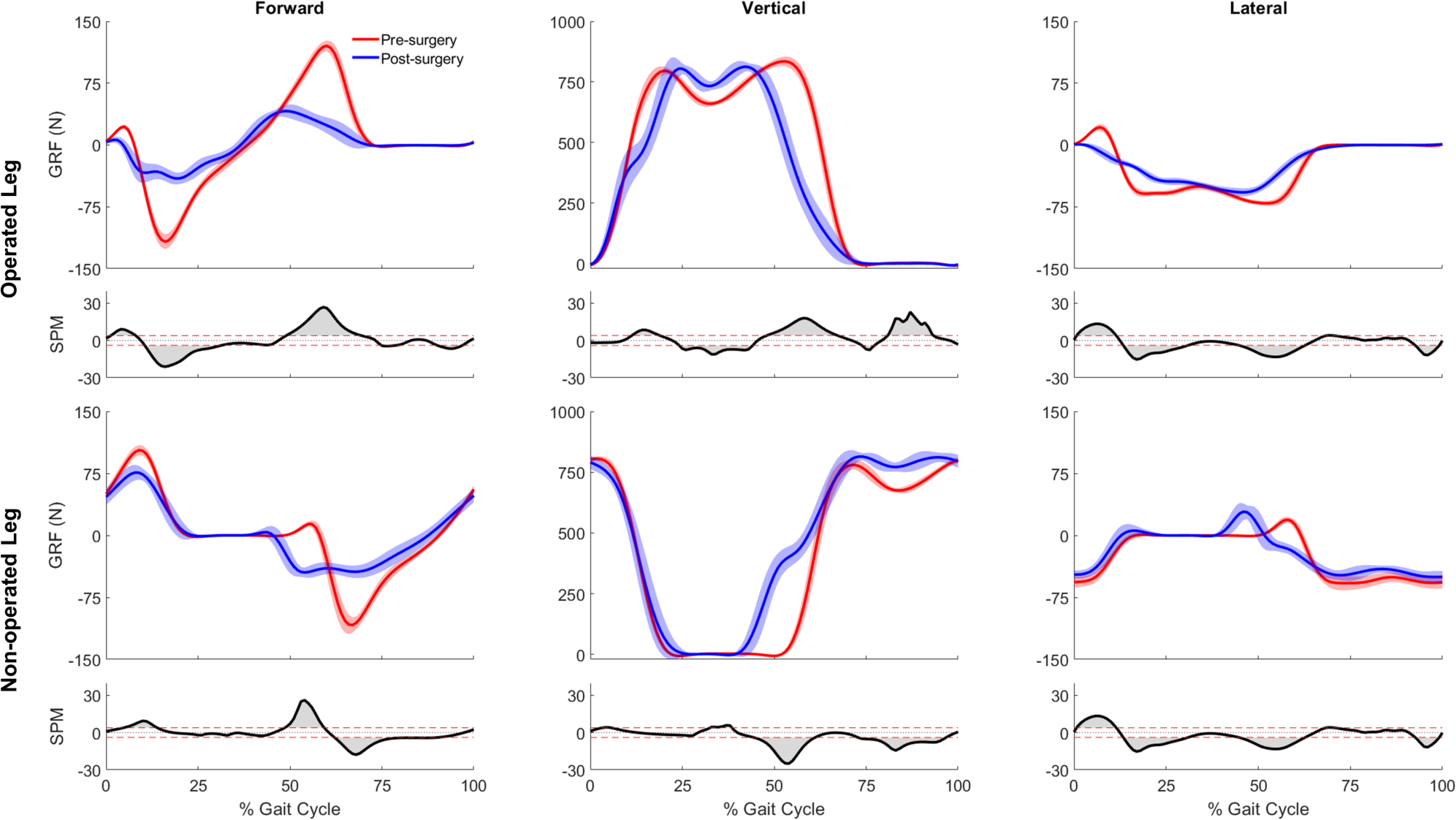
Pre- and post-surgery ground reaction forces (mean ± 1 standard deviation across 10 gait cycles), along with SPM test results (grey shaded area indicates significant difference between pre- and post-surgery data).

For joint motion data, all joint angle trajectories for both the operated and non-operated leg showed significant changes between pre- and post-surgery conditions (*p* < 0.001). For the operated leg following surgery (Fig. 3), the hip was more extended and externally rotated over most of the gait cycle, the knee was less flexed over most of the gait cycle, and the ankle was less dorsiflexed and the subtalar joint less inverted during stance phase. In contrast, for the non-operated leg following surgery (Fig. 3), the hip was more extended only during the first half of stance phase and more internally rotated over most of the gait cycle, the knee was more flexed during stance phase but less flexed during swing phase, the ankle was more dorsiflexed during the first half of stance phase, and subtalar joint exhibited reduced inversion in the middle of stance phase. For the pelvis and trunk (Fig. 4), joint rotations generally became more asymmetric following surgery, with pelvis tilt losing its stereotypical double humped pattern, pelvis list showing a drop toward the non-operated side, pelvis rotation exhibiting an offset that moved the operated hip forward, lumbar extension demonstrating an increase in forward tilt, and lumbar bending increasing toward the operated side.

**Figure 3.**
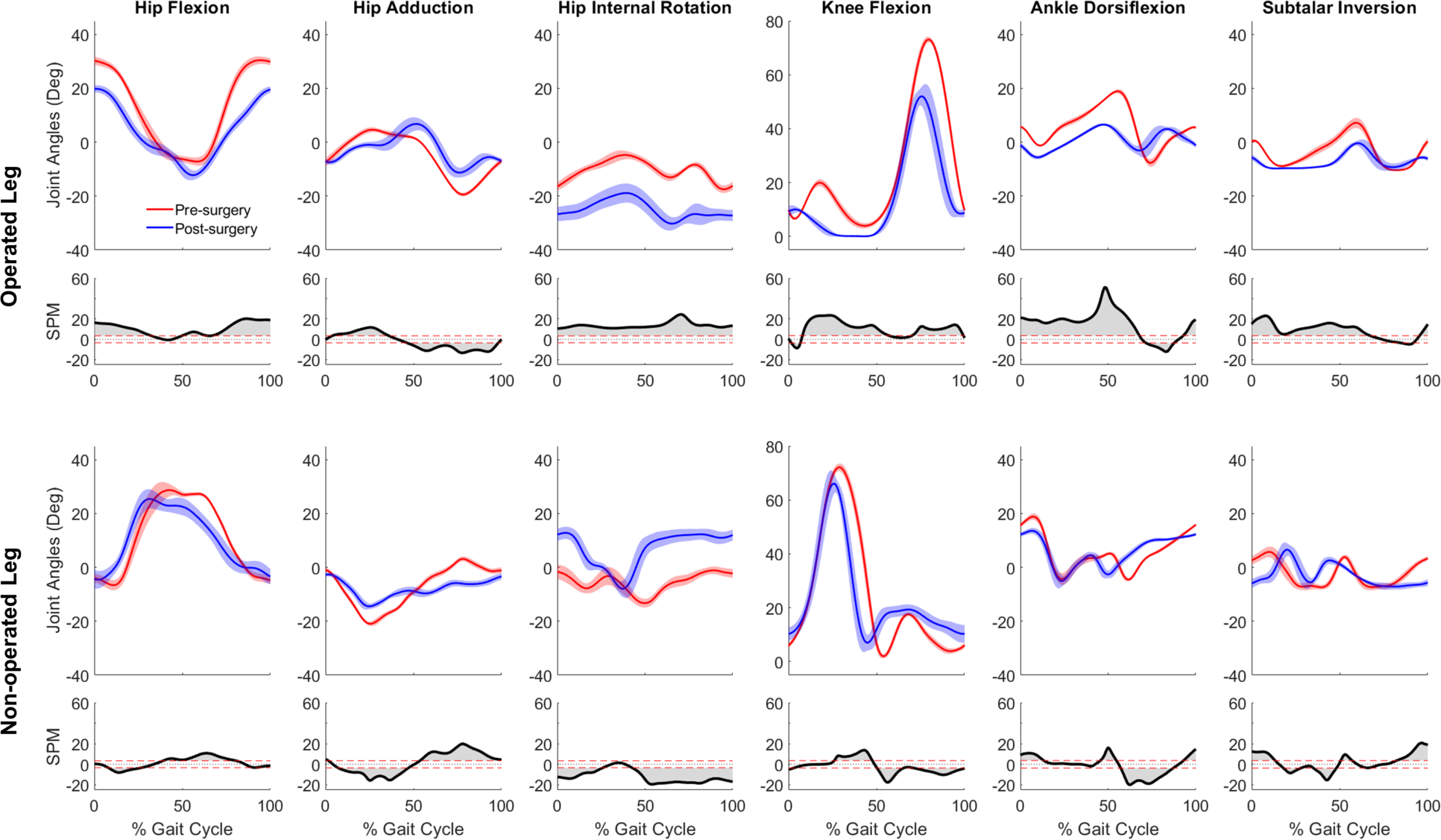
Pre- and post-surgery joint motions (mean ± 1 standard deviation across 10 gait cycles) for all lower extremity joints, along with SPM test results (grey shaded area indicates significant difference between pre- and post-surgery data)

**Figure 4.**
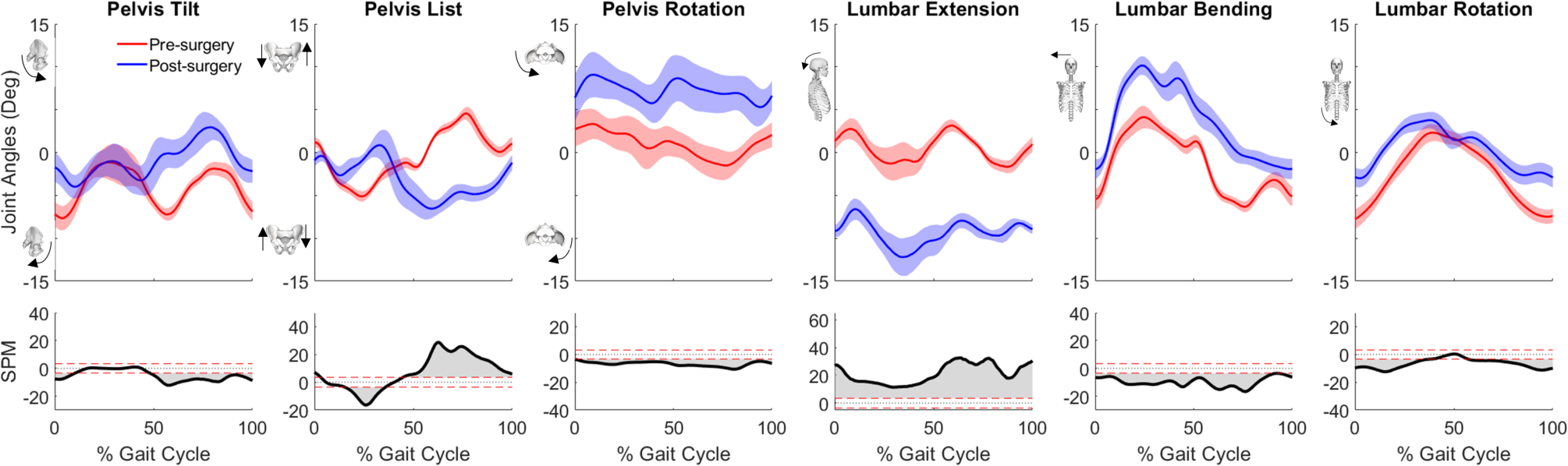
Pre- and post-surgery joint motions (mean ± 1 standard deviation across 10 gait cycles) for pelvis and lumbosacral orientations, along with SPM test results (grey shaded area indicates significant difference between pre- and post-surgery data)

For joint moment data (Fig. 5), more pronounced changes (*p* < 0.001 over larger portions of the gait cycle) occurred in the operated leg than in the non-operated leg following surgery. For the operated leg during stance phase, the hip flexion and abduction moments decreased, the knee extension moment turned into a flexion moment, and the subtalar inversion moment increased. Changes during swing phase were minimal. In contrast, for the non-operated leg, joint moment changes were minimal following surgery, with scattered statistically significant differences occurring at various points throughout the gait cycle.

**Figure 5.**
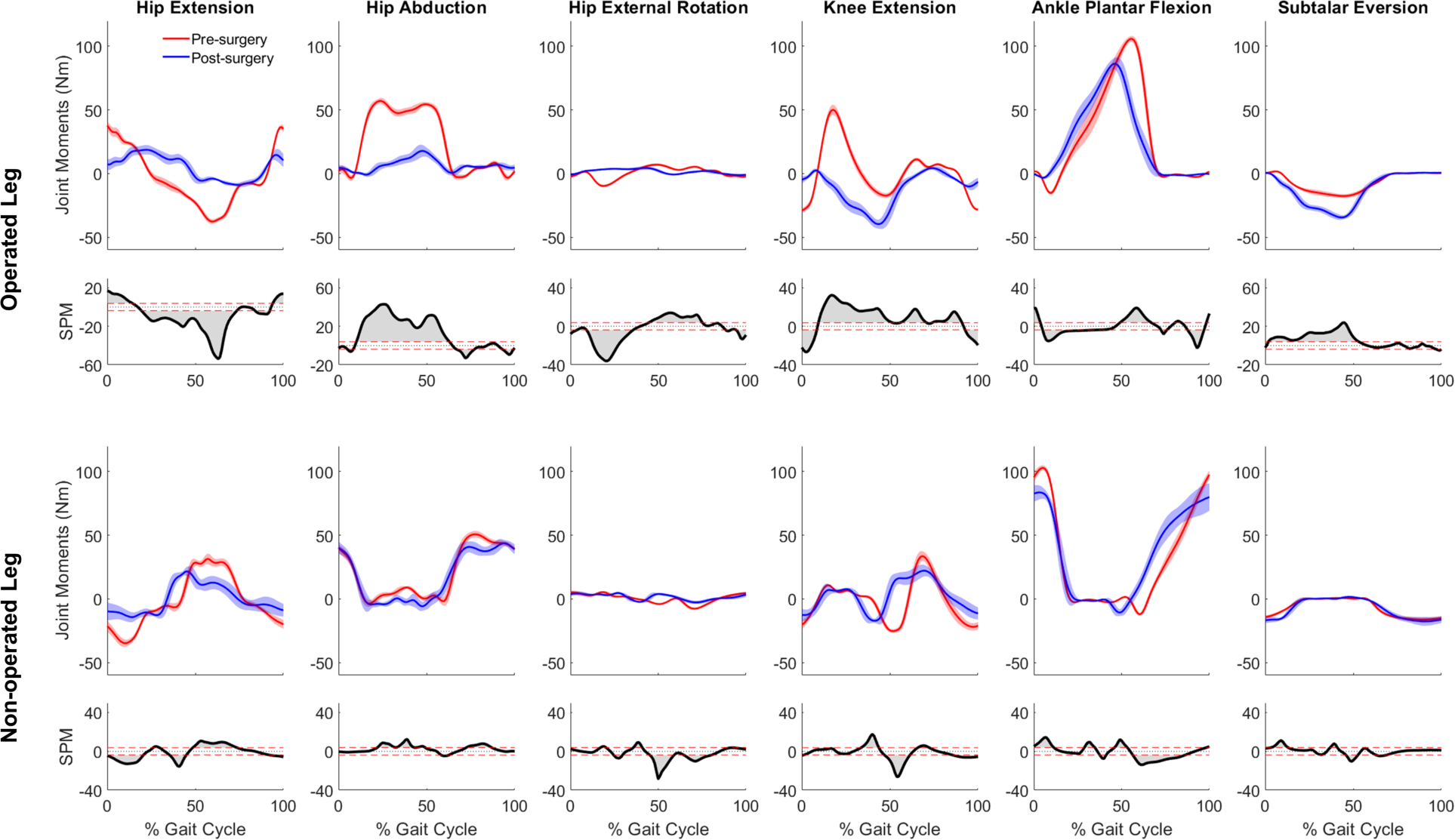
Pre- and post-surgery joint moments (mean ± standard deviation across 10 gait cycles) for all lower extremity joints, along with SPM test results (grey shaded area indicates significant difference between pre- and post-surgery data)

### 3.2 Muscle-tendon Model Parameter Changes

Optimal muscle fiber length and tendon slack length values found by EMG-driven neuromusculo-skeletal model calibration showed relatively small differences on average between pre- and post-surgery conditions (Fig. 7). For optimal fiber length, the changes ranged from −0.019 to 0.070 m (mean 0.018 m and median 0.013 m) for the operated leg, and from −0.050 to 0.044 m (mean −0.003 m and median −0.0005 m) for the non-operated leg. In terms of percentages with respect to pre-surgery values, these changes ranged from −24% to 40% (mean 14% and median 13%) for the operated leg, and from −28% to 132% (mean 2% and median −1%) for the non-operated leg. For tendon slack length, the changes ranged from −0.051 to 0.013 m (mean −0.011 m and median −0.008 m) for the operated leg, and from −0.082 to 0.027 m (mean −0.013 m and median −0.005 m) for the non-operated leg. In terms of percentages with respect to pre-surgery values, the changes ranged from −40% to 5% (mean −9% and median −3%) for the operated leg, and from −70% to 43% (mean −11% and median −5%) for the non-operated leg.

### 3.3 Neural Control Changes

When post-surgery muscle synergies were compared to pre-surgery muscle synergies, stronger similarities were found between synergy excitations or activations than between synergy vectors (Table 2). The cosine similarity values between the pre- and post-surgery synergy controls were generally higher than those between the synergy vectors for both the operated and non-operated leg. When the pre-surgery synergy controls were shifted to maximize cosine similarity (Supplementary Figure 5), the cosine similarity values between the optimally shifted pre-surgery and post-surgery synergy controls were close to one (Table 2), indicating extremely strong similarity. The shifts required to achieve maximum cosine similarity (Supplementary Table 3) appeared to decrease with an increasing number of synergies. Figure 6 illustrates the comparison between paired pre- and post-surgery muscle synergy quantities and how cosine similarity values were calculated between paired synergy activations and synergy vectors.

**Figure 6.**
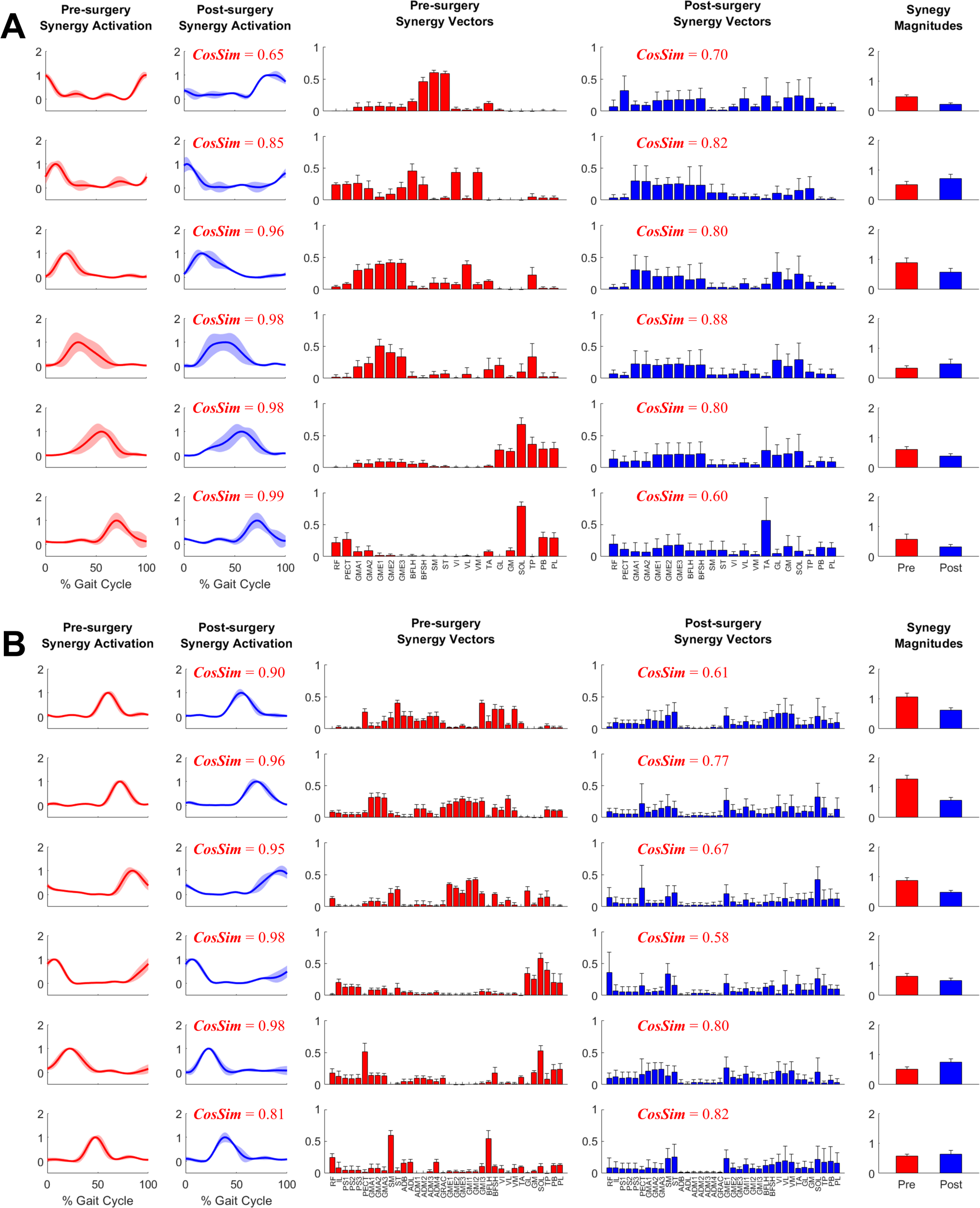
Example plot of pre- and post-surgery muscle synergies (mean ± 1 standard deviation across 10 gait cycles) for A. Operated Leg and B. Non-operated Leg. Cosine similarity was calculated using the mean values for each pair of synergy activations or synergy vectors.

**Table 2:** Comparison of pre- and post-surgery muscle synergies using the mean ± standard deviation of cosine similarity between pre- and post-surgery muscle synergy quantities across synergies. For each pair of pre- and post-surgery muscle synergies identified after sorting and pairing, we calculated cosine similarity between (1) mean muscle synergy control curves and (2) mean synergy vectors. Mean synergy controls and synergy vectors were calculated from all ten pre- or post-surgery gait cycles analyzed. MSA was performed using both muscle excitations and muscle activations to determine potential differences caused by the type of muscle control analyzed.

| Number of Synergies | Muscle Control | Cosine Similarity |  |  |  |  |  |
| --- | --- | --- | --- | --- | --- | --- | --- |
|  |  | Non-operated Leg |  |  | Operated Leg |  |  |
|  |  | (1)<br>Synergy controls | (2)<br>Shifted controls | (3)<br>Synergy vectors | (1)<br>Synergy controls | (2)<br>Shifted controls | (3)<br>Synergy vectors |
| 4 | Excitation | 0.77±0.18 | 0.95±0.03 | 0.71±0.11 | 0.90±0.08 | 0.95±0.03 | 0.75±0.15 |
| | Activation | 0.84±0.11 | 0.94±0.02 | 0.78±0.05 | 0.74±0.25 | 0.98±0.01 | 0.68±0.20 |
| 5 | Excitation | 0.90±0.05 | 0.95±0.05 | 0.83±0.09 | 0.86±0.12 | 0.96±0.05 | 0.73±0.10 |
| | Activation | 0.89±0.05 | 0.95±0.02 | 0.75±0.09 | 0.84±0.19 | 0.98±0.02 | 0.72±0.14 |
| 6 | Excitation | 0.94±0.03 | 0.97±0.02 | 0.79±0.05 | 0.87±0.05 | 0.95±0.05 | 0.76±0.08 |
| | Activation | 0.93±0.06 | 0.98±0.01 | 0.71±0.10 | 0.90±0.13 | 0.97±0.02 | 0.77±0.10 |
| 7 | Excitation | 0.76±0.18 | 0.96±0.02 | 0.78±0.04 | 0.91±0.12 | 0.97±0.02 | 0.77±0.09 |
| | Activation | 0.94±0.07 | 0.98±0.01 | 0.81±0.08 | 0.96±0.03 | 0.97±0.03 | 0.82±0.05 |

### 3.4 Post-surgery Muscle Control Predictions

When pre-surgery muscle synergy quantities were used to fit post-surgery muscle controls, better fits were obtained using pre-surgery synergy controls rather than pre-surgery synergy vectors (Fig. 8). Both the Fixed Synergy Control and Shifted Synergy Control method achieved higher VAF than did the Fixed Synergy Vector method for fitting post-surgery muscle excitations and activations with any number of synergies between 4 and 7 (*p* < 0.001). The Shifted Synergy Control method applied to muscle activations achieved higher VAF than did the corresponding Fixed Synergy Control method using 5, 6, and 7 synergies for the non-operated leg (*p* < 0.05) and 4, 5, and 6 synergies for the operated leg (*p* < 0.05). The Fixed Synergy Control method performed significantly better when applied to muscle activations rather than muscle excitations except for the operated leg with 7 synergies (*p* = 0.0591). The performance of both the Fixed Synergy Control and Shifted Synergy Controls methods applied to muscle activations generally improved with increasing number of synergies, except when the number of synergies increased from 6 to 7 for the operated leg (*p* = 0.1065).

**Figure 7.**
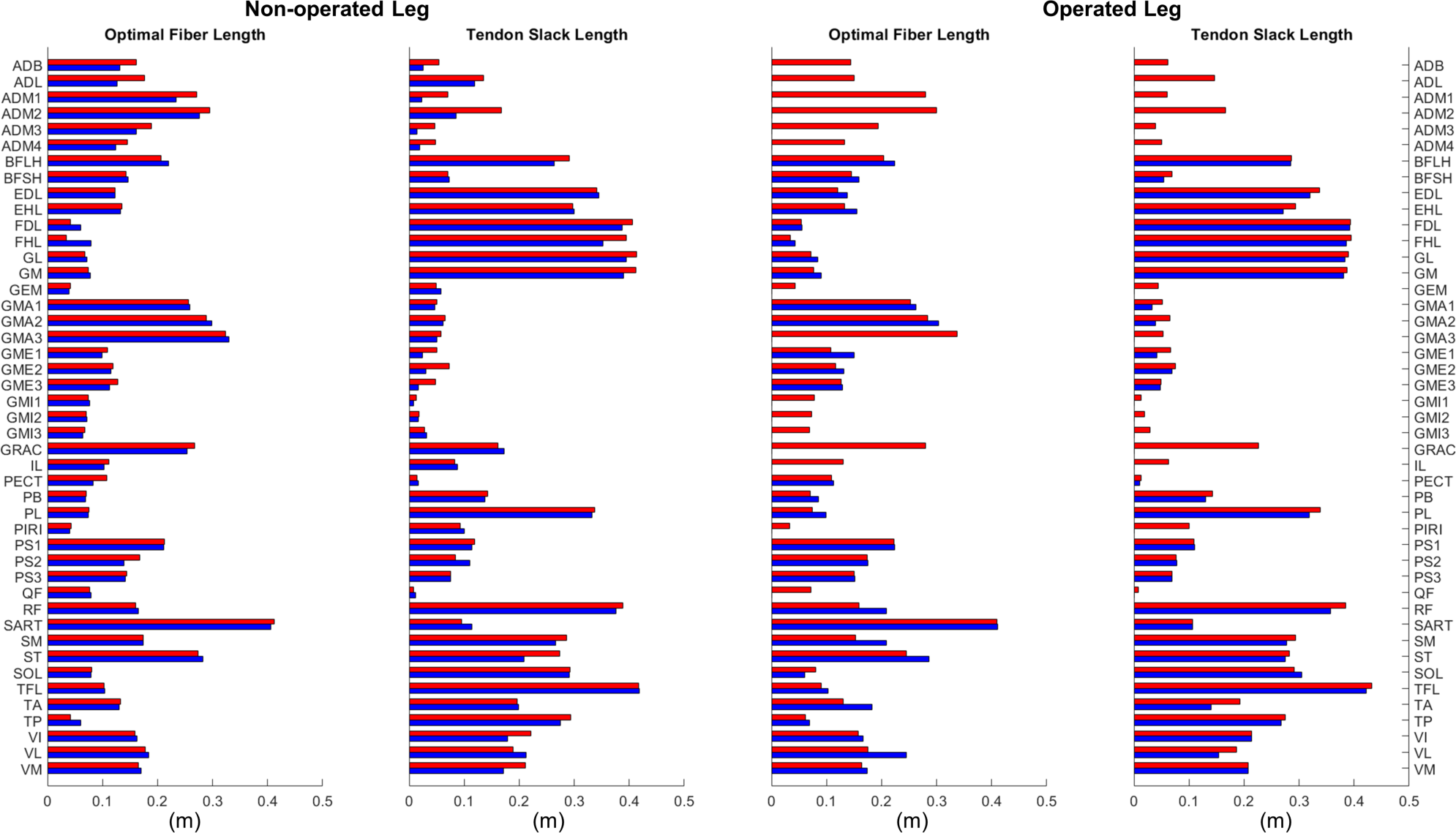
Optimal muscle fiber length and tendon slack length values for lower extremity muscles in the personalized pre-surgery (red) and post-surgery (blue) neuromusculoskeletal models.

**Figure 8.**
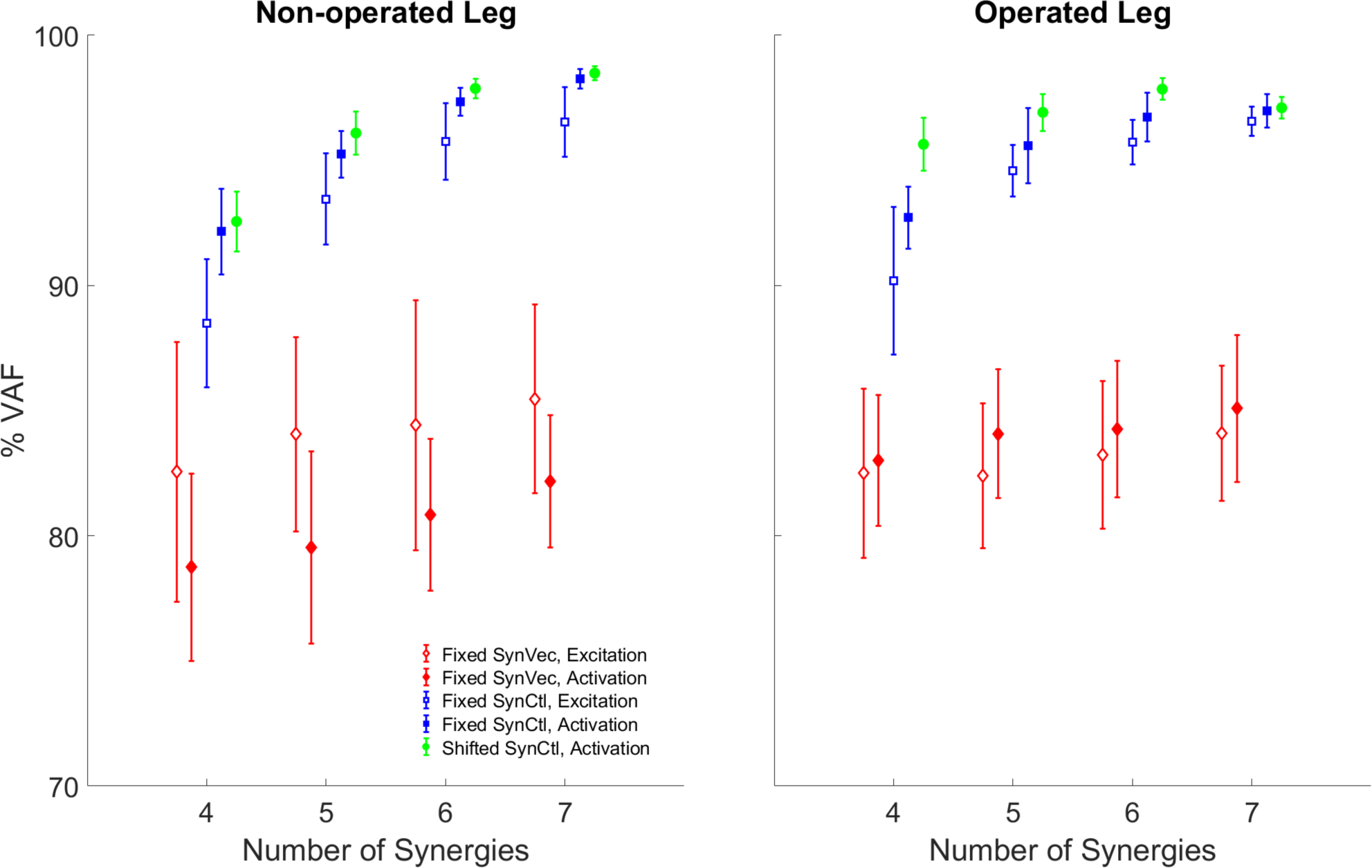
Variability accounted for (VAF) by reconstructed post-surgery muscle controls using each method: FixedSynVec (Fixed Synergy Vector, red), FixedSynCtl (Fixed Synergy Control, blue), and ShiftedSynCtl (Shifted Synergy Control, green). Each marker with error bars indicates mean ± 1 standard deviation of VAF values from 10 gait cycles. Open markers indicate reconstruction of muscle excitations while filled markers indicate reconstruction of muscle activations.

## 4 Discussion

This study analyzed changes in walking function and neural control for a single pelvic sarcoma patient after internal hemipelvectomy surgery with custom prosthesis reconstruction. Personalized neuromusculoskeletal computational models representing the subject before and after surgery were developed to quantify changes in biomechanical quantities, neuromusculoskeletal model parameter values, and neural control quantities due to the surgery. Muscle synergy analyses were performed on the experimental muscle excitations and associated muscle activations for lower extremity muscles with experimental EMG data to quantify changes in neural control. Three methods for predicting post-surgery muscle controls using pre-surgery muscle synergy information were evaluated. Our findings suggest that for the subject analyzed in this study, post-surgery walking function differs substantially from pre-surgery walking function despite a post-surgery gait pattern that does not appear visually to be largely abnormal. These quantified differences could provide targets for personalized rehabilitation efforts for this subject. Furthermore, our findings suggest that the Fixed and Shifted Synergy Control methods reconstructed post-surgery muscle controls better than did the Fixed Synergy Vector method. The total VAF of reconstructed post-surgery muscle activations reached 95% using only five synergies for the Shifted Synergy Control method and six synergies for the Fixed Synergy Control method. Consequently, future studies that seek to use personalized computational neuromusculoskeletal models to predict post-surgery walking function from pre-surgery walking data and the planned surgical decisions should explore using the Fixed or Shifted Synergy Control method to model the subject’s post-surgery neural control strategy.

Compared to previous studies, the present study collected more extensive walking data, with the data being available both before and after surgery. In part because custom prosthesis reconstruction has become a viable option only recently, few studies have performed instrumented gait analyses of this patient population (Akiyama et al., 2016; Wingrave and Jarvis, 2019; Valente et al., 2022). As the emphasis of these previous studies was on evaluating post-surgery walking function, pre-surgery walking data were not available from the same patients. No previous study reported ground reaction data for all three components of ground reaction force or joint moment data for all lower body joints. Only one study reported EMG data collected from six muscles in the operated leg. Thus, while limited to a single subject, the experimental walking data collected for the present study will be a valuable resource that other researchers can use to perform their own investigations of how internal hemipelvectomy surgery with custom prosthesis reconstruction affects post-surgery walking function.

While the extensive walking data collected for this study allowed for a detailed analysis of walking function and neural control changes between pre- and post-surgery conditions, the ultimate goal for these data is to facilitate the development of personalized neuromusculoskeletal modeling methods that can predict a subject’s post-surgery walking function reliably given the subject’s pre-surgery walking data and the surgical decisions being planned by the orthopedic oncologist. Calibration of personalized neuromusculoskeletal models currently requires extensive walking data, including motion capture, ground reaction, and EMG data. Furthermore, these data must be available before and after surgery so that the ability of the subject’s pre-surgery model to predict post-surgery walking function can be evaluated. One of the key challenges in predicting post-surgery walking function is that the subject’s post-surgery neural control strategy must be predicted at the same time. Thus, the availability of extensive pre- and post-surgery walking data from the same subject provides a unique opportunity to develop a hypothesis for how to model a subject’s post-surgery neural control strategy starting from the subject’s pre-surgery walking data.

Despite a post-surgery gait pattern that visually appeared to be only mildly abnormal, the subject’s gait pattern was substantially altered by the surgery. First and foremost, the subject decreased his self-selected walking speed from 1.0 m/s pre-surgery to 0.5 m/s post-surgery. Since self-selected walking speed was cut in half, it is not surprising that the subject exerted less propulsive and braking force following surgery (Fig. 2). Second, the subject exhibited a shortened swing phase for the non-operated leg post-surgery, likely a compensatory mechanism to reduce operated leg single-limb support time. Third, the subject exhibited post-surgery compensatory gait changes consistent with a Trendelenburg-Duchenne gait pattern (Kiernan et al., 2018). These changes included pelvis drop to the non-operated side coupled with lumber bending to the operated side during operated side single leg support (Fig. 4), which is consistent with impaired hip abductor function (Supplementary Table 1) and a significantly reduced hip abduction moment in the operated leg following surgery (Fig. 5). Fourth, the subject also exhibited post-surgery compensatory gait changes consistent with stiff knee gait (Fujita et al., 2022). The subject exhibited a hyperextended knee from mid to late stance phase (Fig. 3A), producing a knee flexion moment rather than the expected knee extension moment. This atypical knee moment counteracted the external moment caused by the ground reaction force vector passing in front of rather than behind the knee, which in turn was caused by significant forward trunk lean following surgery (Fig. 4). Though knee hyperextension and greater anterior trunk lean have been observed in individuals with quadriceps muscle weakness (Siegel et al., 2007; Sato and Maitland, 2008), the subject’s vastii muscles were not touched by the surgery, while his rectus femoris muscle was detached at its origin but later re-attached. Thus, while quadriceps muscle weakness seems unlikely, the subject’s post-surgery EMG data suggests that quadriceps muscle activation impairment may have occurred (Supplementary Fig. 3A) for reasons that remain unclear.

To support our ultimate goal of developing a computational methodology that can predict a subject’s post-surgery walking function from his pre-surgery walking data and the planned surgical decisions, we calculated muscle synergies that were not only electromyographically consistent but also kinetically consistent. In nearly all published studies, muscle synergies are calculated from experimental EMG data alone, making them only electromyographically consistent. If the calculated muscle synergies were input into a neuromusculoskeletal model of the subject, along with experimental joint kinematic data, the model would not produce the subject’s experimental joint moments and thus would be kinetically inconsistent. In contrast, muscle synergies in our study were calculated from experimental EMG, joint kinematic, and joint moment data via calibrated EMG-driven musculoskeletal models, making them not only electromyographically consistent but also kinetically consistent. Thus, our calculated muscle synergies should provide an excellent starting point for future predictive simulations of post-surgery walking function.

Two observations support the conclusion that the Fixed or Shifted Synergy Control method is a better choice than the Fixed Synergy Vector method for predicting post-surgery muscle controls within a predictive simulation of post-surgery walking function. First, the largest pre-to post-surgery changes in neural control, as quantified using muscle synergies, occurred in the subject’s synergy vectors rather than his synergy excitations or activations. The cosine similarity metric showed that there was higher similarity between pre- and post-surgery synergy controls than between pre- and post-surgery synergy vectors (Table 2). The fact that the subject’s synergy vectors exhibited substantial pre-to post-surgery changes for both legs may be the first published evidence that an individual with healthy neural control is able to change his coordination strategy in both legs in response to the loss of a significant number of muscles in one leg. Second, the Fixed and Shifted Synergy Control methods worked much better than did the Fixed Synergy Vector method for reconstructing post-surgery muscle controls using pre-surgery synergy information (Fig. 8). The Fixed Synergy Vector method assumes that the coordination strategy for lower extremity muscles stays more or less the same follow surgery. However, this method was incapable of accurately reconstructing post-surgery muscle controls, whereas the Fixed and Shifted Synergy Control methods achieved highly accurate post-surgery reconstructions.

Another important question for predicting post-surgery changes in neural control is whether the neural control model should represent muscle excitations or muscle activations. When post-surgery muscle controls were reconstructed using the Fixed or Shifted Synergy Control method, the highest reconstruction accuracy for both legs was achieved when the reconstruction process was applied to muscle activations rather than muscle excitations (Fig. 8). This difference may be related to the fact that muscle activations are “closer” to biomechanical function than are muscle excitations, which must be time delayed and passed through activation dynamics to be transformed into muscle activations. Electromechanical delay, which was roughly constant for all muscles, would equate to different percentages of the gait cycle for different walking speeds. The average calibrated electromechanical delay for all muscles was 83.7 milliseconds. This delay equated to an average 7.6% of the pre-surgery gait cycle (1.0 m/s) but only 5.6% of the post-surgery gait cycle (0.5 m/s). Thus, from the standpoint of a normalized gait cycle, one might expect pre- and post-surgery muscle activations to be better synchronized than pre- and post-surgery muscle excitations.

While reconstruction of post-surgery muscle controls using the Fixed or Shifted Synergy Control method generally improved with an increased number of muscle synergies, it is important to consider how many synergies should be used. Given our ultimate goal of generating predictive simulations of post-surgery walking function, the lowest number of synergies should be chosen that can reconstruct post-surgery muscle controls accurately. An accurate reconstruction capable of controlling a predictive walking simulation requires a VAF value higher than 95% overall and 85% for individual muscles (Meyer et al., 2016). For the Fixed Synergy Control method with five synergies, the mean overall VAF value for muscle activation reconstruction was 95.2% for the non-operated leg and 95.6% for the operated leg, though overall VAF values for several gait cycles failed to reach the 95% threshold (Fig. 8). Using six synergies, the overall and individual muscle VAF values for reconstructed muscle activations both exceeded the previous requirements. When seven synergies were used, the VAF improvements started to diminish, especially for the operated leg (*p* = 0.1065). For the Shifted Synergy Control method, only five synergies were needed to achieve the desired overall and individual muscle VAF criteria for both the non-operated and operated leg, representing a less expensive computational option for predictive simulations. Thus, for future predictive simulation studies of post-surgery walking function, it would be difficult to justify using more than five or six synergies per leg.

In addition to changes in neural control, quantifying changes in muscle-tendon model parameters value is helpful for future studies that perform predictive simulations of post-surgery walking function. These parameter values, namely optimal muscle fiber length and tendon slack length, are crucial for modeling the force-generating characteristics of the patient’s lower extremity muscles. Based on a comparison of the patient’s pre- and post-surgery calibrated neuromusculoskeletal models, changes in these parameter values were generally small despite some outliers (Fig. 7). These outliers generally occurred for two types of muscles. The first type included muscles without experimental EMG data (Table 1). For example, flexor hallucis longus (FHL) in the non-operated leg had its optimal muscle fiber length increased by 0.045 m or 132%. Since the excitation of such muscles had to be estimated using synergy extrapolation, associated muscle-tendon model parameter values tended to be less accurate than for muscles with experimental EMG data. The second type included muscles that were somehow affected by the surgery. For example, vastus lateralis (VL) in the operated leg had its optimal fiber length increased by 0.070 m or 40%. As noted previously, the patient avoided using this muscle following surgery for reasons that remain unclear. Parameter tuning for such muscles during model calibration was possibly more aggressive to overcome the deficits in activation. With removal of these two types of outliers, changes in these model parameter values were much smaller. The median change in optimal muscle fiber length was 0.013 and −0.0005 m for the operated and non-operated leg, respectively. The median change in tendon slack length was −0.008 and −0.005 m for the two legs. The median values of percent change were also small (13%, −1%, −3%, and −5%, respectively). Thus, for future studies that seek to predict post-surgery walking function starting from a pre-surgery walking model, pre-surgery muscle-tendon model parameter values should provide a reasonable approximation.

This study possesses several limitations related to data generalizability and computational methodology. First, this study analyzed data collected from a single pelvic sarcoma patient. Given the significant heterogeneity between patients with this pathology, it is unknown whether the conclusions drawn from the present study can be generalized to other patients with a pelvic sarcoma. More data collected from more patients are needed to determine generalizability of these results. The number of patients with a pelvic sarcoma is limited, plus it is challenging to find patients who are willing to come in for gait testing prior to surgery and after plateau in recovery. However, the findings of this study can still be used as the foundation for the first predictive simulations of post-surgery walking for this patient population. Second, during calibration of post-surgery EMG-driven models, the peak isometric force values of the remaining hip muscles in the operated leg were set at 100% of their pre-surgery values and were not modified despite the possibility that some muscles may have been weakened by the surgery. Use of pre-surgery peak isometric force values could lead to underestimation of activations for certain hip muscles, especially those that were detached and later reattached. Third, although measured excitations were available for the majority of lower extremity muscles (Table 1), excitations of multiple unmeasured muscles still needed to be estimated using synergy extrapolation (Bianco et al., 2018; Ao et al., 2020, 2022). While this method of estimating unmeasured muscle excitations has been shown to be reliable for a small number of muscles, its effectiveness when extended to multiple muscles has yet to be verified. Fourth, the methods investigated for reconstructing post-surgery muscle controls from pre-surgery muscle synergy information were evaluated using only experimental muscle excitations and activations. The reliability of these methods for predicting the activations of muscles without experimental EMG data is unknown.

In conclusion, this study performed extensive analyses of walking function and neural control changes for a single pelvic sarcoma patient following internal hemipelvectomy surgery with custom pelvic prosthesis reconstruction. The patient exhibited substantial changes in his post-surgery walking function when quantified using experimental data, despite only minor abnormalities being observed visually. The observed changes could potentially provide valuable information for designing a personalized rehabilitation protocol for this patient. The patient also exhibited substantial changes in the coordination of his lower extremity muscles in both legs, as evidenced by pre-to post-surgery changes in his synergy vectors. Consequently, when pre-surgery muscle synergy information was used to reconstruct post-surgery muscle activations, the Fixed and Shifted Synergy Control methods that used pre-surgery synergy activations but found new synergy vectors produced the most accurate reconstructions (>95% VAF) and required only five or six synergies. Consequently, we recommend that future computational studies that seek to predict post-surgery walking function for this patient population start by using the patient’s pre-surgery synergy activations, thereby greatly simplifying the process of predicting the patient’s post-surgery neural control strategy.

## 6 Author Contributions

GL: Conceptualization, Formal analysis, Investigation, Methodology, Software, Visualization, Writing – original draft; DA: Methodology, Software, Writing – reviewing & editing; MV: Methodology, Software, Writing – reviewing & editing; PZ: Data curation, Writing – reviewing & editing; S-HC Data curation, Writing – reviewing & editing; AP: Resources, Writing – reviewing & editing; VL: Conceptualization, Resources, Writing – reviewing & editing; BF: Conceptualization, Data curation, Funding acquisition, Project administration, Software, Supervision, Writing – original draft, Writing-review & editing.

## Acknowledgments

The authors would like to thank Andrew Baines and Nicholas Dunbar for their assistance with segmenting the patient’s CT scan data.

## 8 Data Availability Statement

None.

---

**Supplementary Figure 1.**
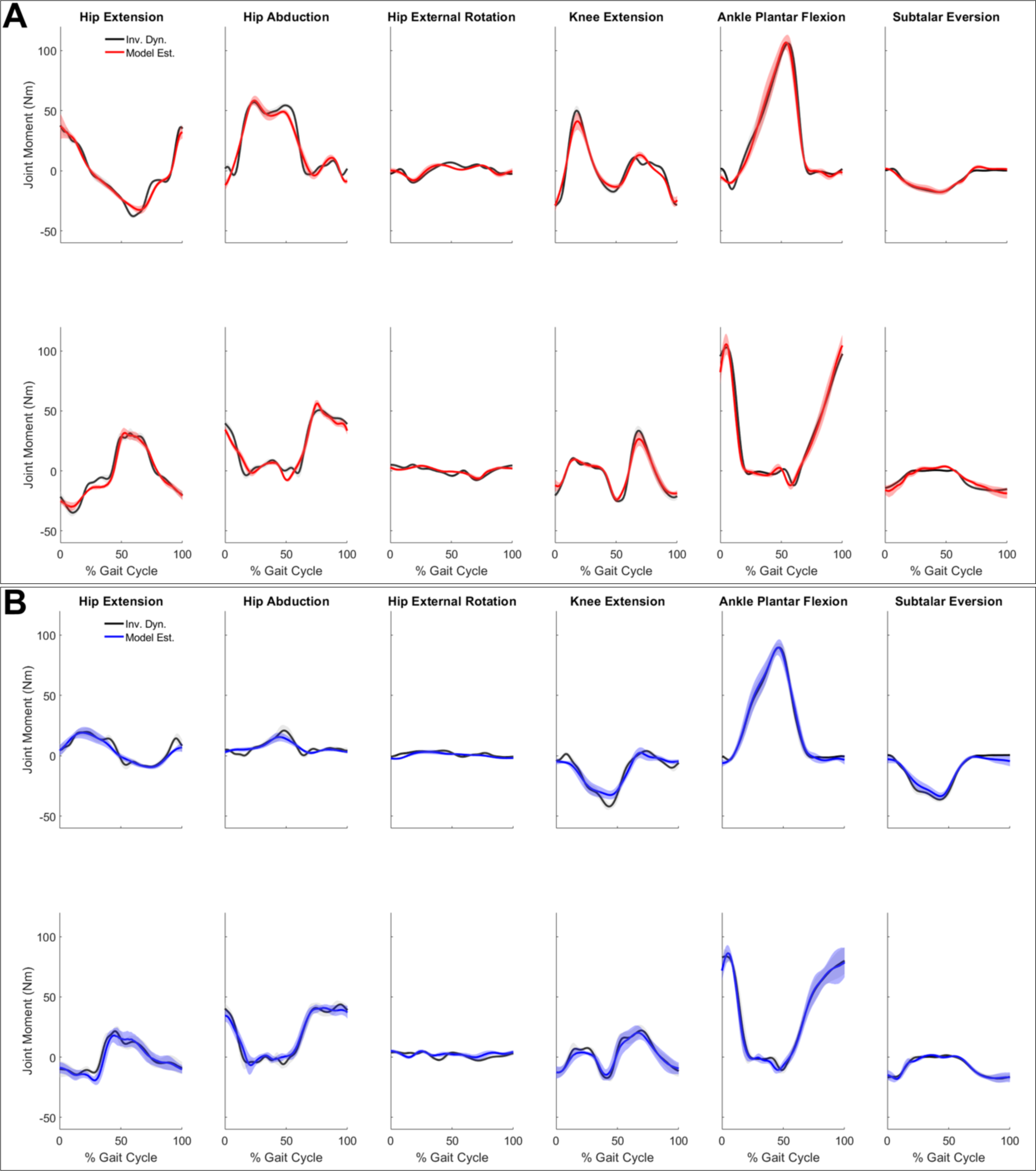
Lower extremity joint moments estimated by the EMG-driven models and calculated from inverse dynamics for **A.** Pre-surgery joint moments: model-estimated (red), inverse dynamics (dark). **B.** Post-surgery joint moments: model-estimated (blue), inverse dynamics (dark). The solid curves represent mean and the shaded areas represent ± 1 standard deviation of joint moments.

**Supplementary Figure 2.**
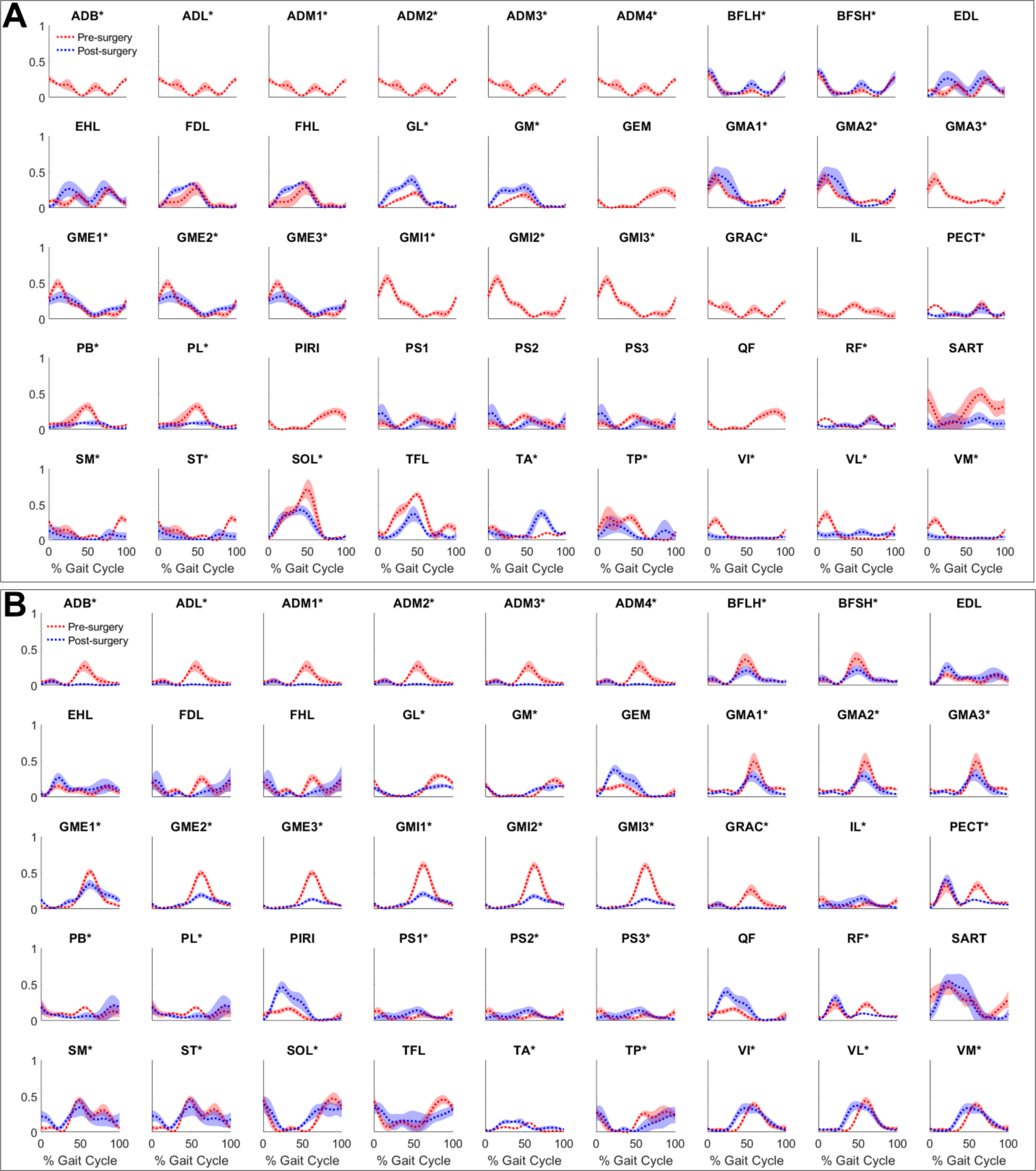
Excitations (mean ± standard deviation across 10 gait cycles) of lower extremity muscles pre-surgery (red) and post-surgery (blue) for **A.** operated leg and **B.** non-operated leg. * indicates EMG data of the muscles were collected during gait trials.

**Supplementary Figure 3.**
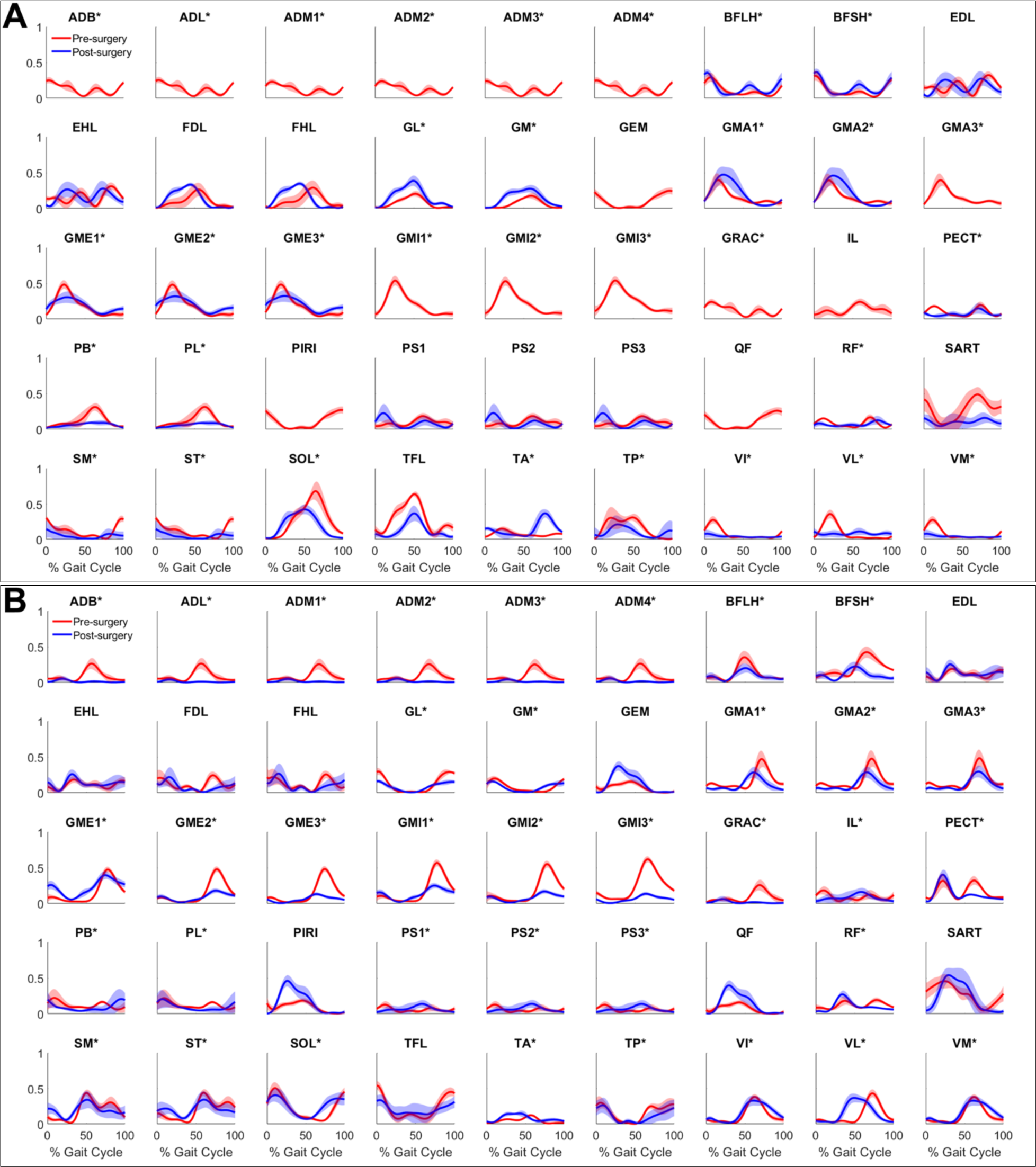
Activations (mean ± standard deviation across 10 gait cycles) of lower extremity muscles pre-surgery (red) and post-surgery (blue) for **A.** operated leg and **B.** non-operated leg. * indicates EMG data of the muscle were collected during gait trials.

**Supplementary Figure 4.**
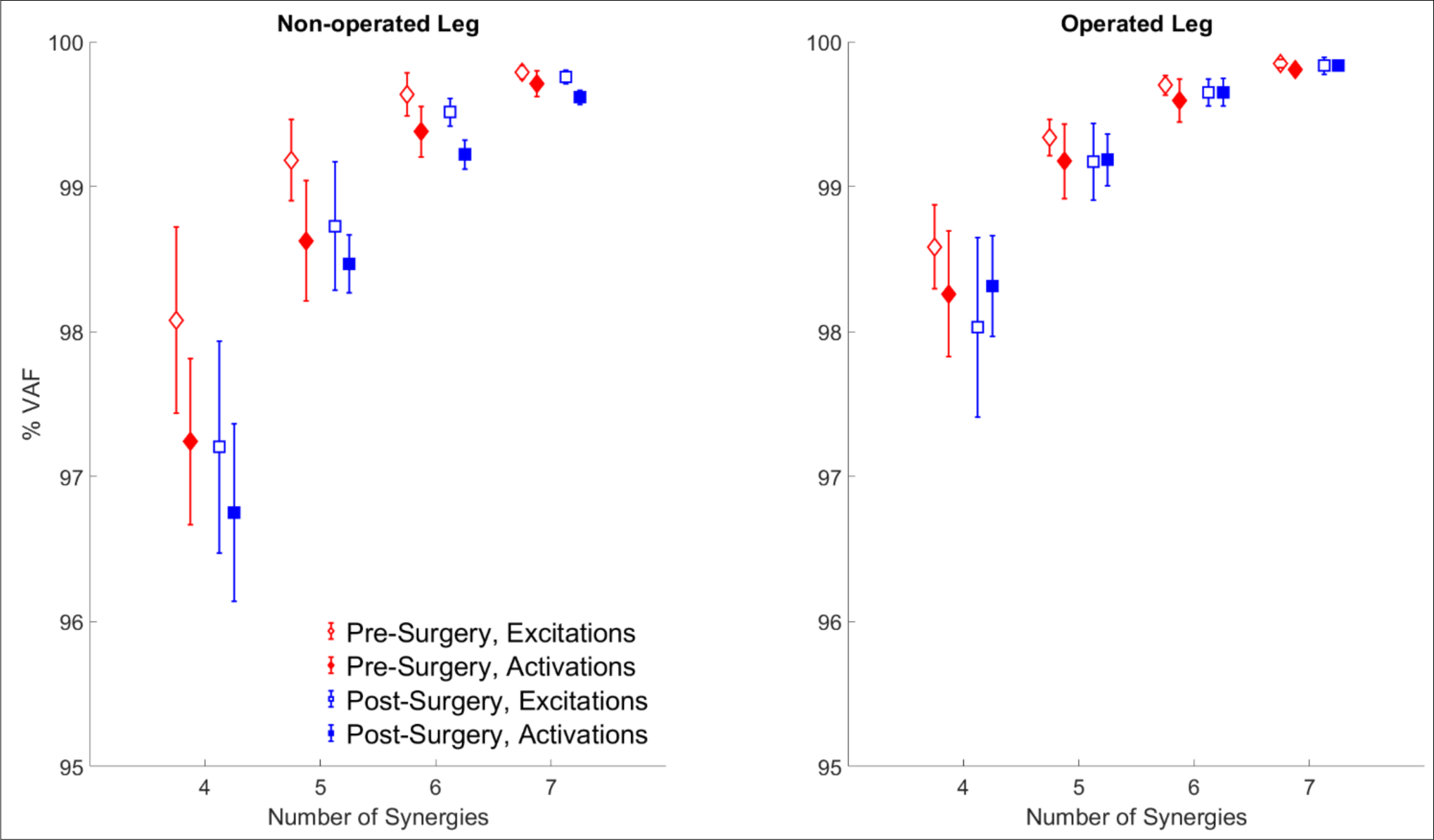
Variability account for, or VAF (mean ± standard deviation across 10 gait cycles) of muscle excitations (empty markers) and activations (filled markers) by the muscle synergies for pre-surgery (red or diamond) and post-surgery (blue or square).

**Supplementary Figure 5.**
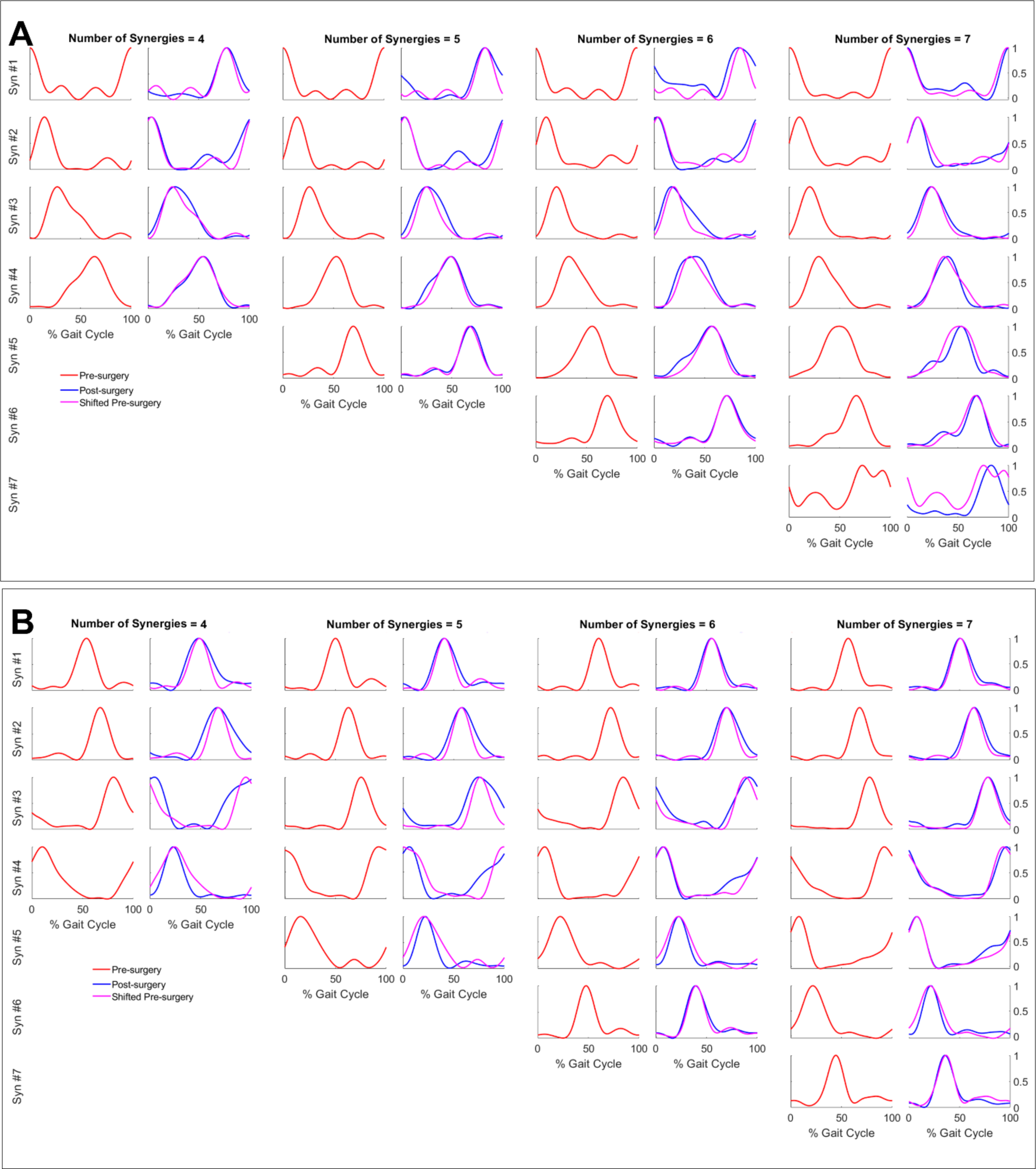
Pre-surgery (red) and post-surgery (blue) synergy activations (mean across 10 gait cycles), as well as the shifted pre-surgery synergy activations (magenta) that had maximum cosine similarity with post-synergy activations for **A.** operated leg and **B.** non-operated leg. See Supplementary Table X for the shifts required to achieve maximum cosine similarity.

**Supplementary Figure 6.**
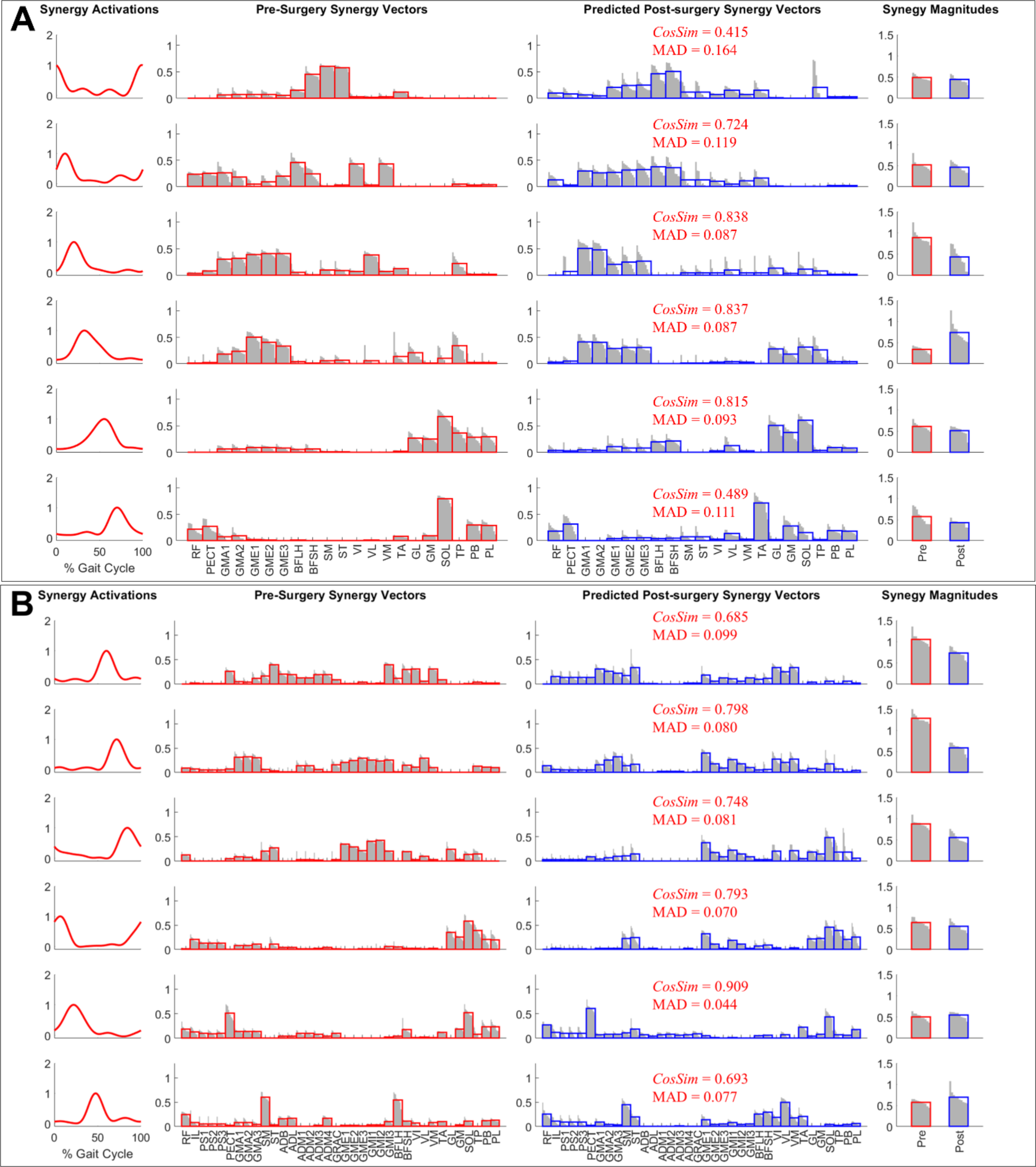
Pre-surgery synergy commands and synergy vectors (red), and synergy vectors required to reconstruct post-surgery muscle activations using Fixed Synergy Control method (blue) for **A.** operated leg and **B.** non-operated leg. The red and blue bars represent the mean values of cycle-specific synergy vector weights (grey). Cosine similarity and mean absolute difference between the two set of synergy vectors were calculated using the mean values (red and blue bars).

**Supplementary Table 1.**
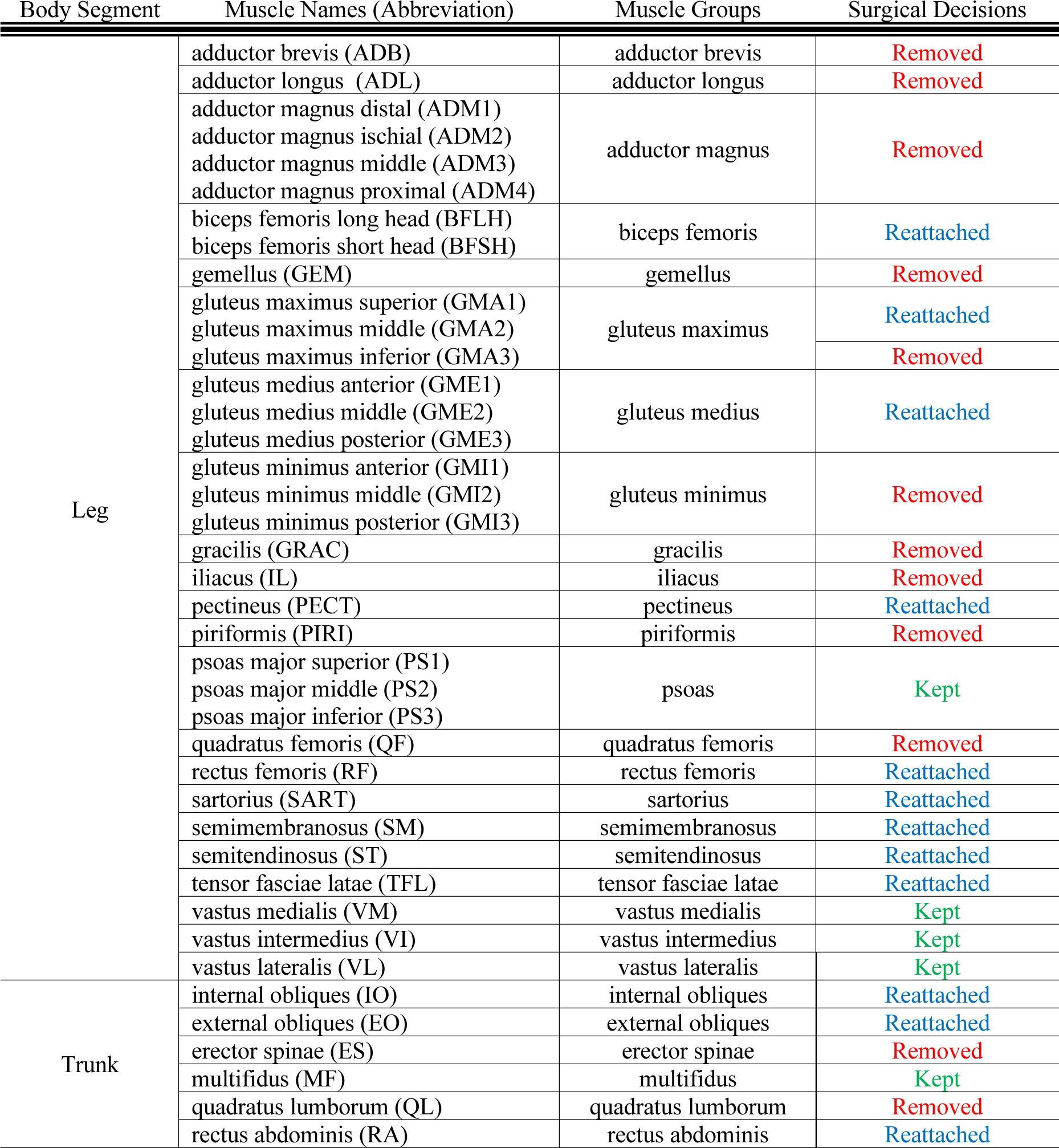
Status of the muscles on the operated hemipelvis (right) following an internal hemipelvectomy surgery.

**Supplementary Table 2.** Mean absolute errors in the joint moments estimated by EMG-driven models. HipFE – hip flexion/extension, HipAA – hip adduction/abduction, HipRot – hip internal/external rotation, KneeFE – knee flexion/extension, AnkleDP – ankle dorsi/plantar flexion, Subtalar IE – subtalar inversion/eversion.

|  |  | Mean Absolute Error in Joint Moment Estimates (Nm) |  |  |  |  |  |
| --- | --- | --- | --- | --- | --- | --- | --- |
|  | Leg | HipFE | HipAA | HipRot | KneeFE | AnkleDP | SubtalarIE |
| Pre-surgery | R | 3.49 | 5.01 | 2.08 | 4.23 | 4.36 | 1.50 |
|  | L | 3.70 | 4.09 | 1.77 | 2.93 | 4.96 | 2.63 |
| Post-surgery | R | 2.95 | 2.54 | 1.64 | 4.09 | 2.94 | 2.84 |
|  | L | 3.23 | 3.94 | 2.31 | 3.60 | 3.57 | 1.37 |

**Supplementary Table 3.** Shift required to achieve maximum cosine similarity between pre- and post-surgery synergy activations, for number of synergies n = 4, 5, 6, and 7. See Supplementary Figure 5 for the synergy activation curves.

|  | Operated Leg |  |  |  |
| --- | --- | --- | --- | --- |
|  | n = 4 | n = 5 | n = 6 | n = 7 |
| Syn #1 | -23 | -17 | -14 | -2 |
| Syn #2 | -11 | -10 | -6 | 1 |
| Syn #3 | -3 | -1 | -1 | 5 |
| Syn #4 | -10 | -4 | 2 | 8 |
| Syn #5 |  | -2 | 3 | 3 |
| Syn #6 |  |  | 2 | 2 |
| Syn #7 |  |  |  | 4 |
| Mean absolute | 11.8 | 6.8 | 4.7 | 3.6 |
|  | Non-operated Leg |  |  |  |
|  | n = 4 | n = 5 | n = 6 | n = 7 |
| Syn #1 | -5 | -9 | -5 | -6 |
| Syn #2 | 0 | -5 | -2 | -4 |
| Syn #3 | 14 | 1 | 4 | 0 |
| Syn #4 | 14 | 7 | 1 | 1 |
| Syn #5 |  | 5 | 0 | 0 |
| Syn #6 |  |  | -8 | -1 |
| Syn #7 |  |  |  | -8 |
| Mean absolute | 8.3 | 5.4 | 3.3 | 2.9 |

